# Effect of thermoneutral housing on MASLD severity, hepatic gene expression, and BAT activation during β3-adrenergic stimulation in mice

**DOI:** 10.1101/2024.10.24.619974

**Authors:** Céline Marie Pauline Martin, Arnaud Polizzi, Valérie Alquier-Bacquié, Marine Huillet, Clémence Rives, Charlène Dauriat, Justine Bruse, Valentine Melin, Claire Naylies, Yannick Lippi, Frédéric Lasserre, JingHong Wan, Rémy Flores-Flores, Justine Bertrand-Michel, Florence Blas-Y-Estrada, Elodie Rousseau-Bacquié, Thierry Levade, Hervé Rémignon, Dominique Langin, Etienne Mouisel, Sophie Lotersztajn, Benoit Chassaing, Laurence Gamet-Payrastre, Hervé Guillou, Sandrine Ellero-Simatos, Anne Fougerat, Nicolas Loiseau

## Abstract

Metabolic dysfunction-associated steatotic liver disease (MASLD), and its more advanced stage metabolic dysfunction-associated steatohepatitis, is the most common chronic liver disease, constituting a major public health issue. No medication is approved for MASLD treatment, and relevant preclinical models are needed to define molecular mechanisms underlying MASLD pathogenesis, and evaluate therapeutic approaches. Here we demonstrated that compared to standard temperature housing, thermoneutral housing aggravated western diet (WD)-induced obesity, diabetes, and steatosis in male mice, which was associated with increased hepatic expression of inflammation- and fibrosis-related genes. Accordingly, compared to standard-housed mice, thermoneutral-housed WD-fed mice developed more severe hepatic inflammation and fibrosis. The liver is the central metabolic organ in whole-body metabolic homeostasis. We used thermoneutrally housed mice with WD-induced MASLD to examine the effect of MASLD during β3- adrenergic stimulation, and found that diet-induced MASLD was associated with defective inter-organ metabolic cross-talk, leading to impaired brown adipose tissue activation.

**Highlights:**

- Thermoneutral housing promotes WD-induced obesity and MASLD in mice
- Thermoneutral housing fosters WD-induced change in gene expression
- Thermoneutral housing fosters hepatic inflammation and fibrosis
- MASLD is associated with defective BAT response to β3-adrenergic stimulation

## INTRODUCTION

Metabolic dysfunction-associated steatotic liver disease (MASLD) is the most common chronic hepatic liver disease. It comprises several pathologies—ranging from steatosis, which is benign and reversible, to the more severe steatohepatitis, which is characterized by inflammation, hepatocyte damage, and progressive fibrosis, and is a predisposing factor for cirrhosis and hepatocellular carcinoma (Younossi et al., 2019; Zhou et al., 2024). A hallmark of MASLD pathogenesis is hepatic triglyceride accumulation. Free fatty acids release from adipose tissue lipolysis is the primary mechanism contributing to lipid overload within hepatocytes, followed by increased hepatic fatty acid synthesis through *de novo* lipogenesis and, to a lesser extent, dietary fat intake (Donnelly et al., 2005; Fabbrini et al., 2008). MASLD is closely associated with metabolic dysfunctions— including insulin resistance, type 2 diabetes, and obesity—hence the recent name change from non-alcoholic fatty liver disease (NAFLD) to MASLD (Eslam et al., 2020a, 2020b; Rinella et al., 2023). In obese insulin-resistant individuals, insulin cannot suppress adipose tissue lipolysis, and the released free fatty acids reach the liver. Quantitative lipidomic analysis of both serum and liver has demonstrated that lipids derived from adipocyte lipolysis drive profound changes of lipid remodeling in the mouse liver (Zhang et al., 2024). Additionally, adipose tissue secretes cytokines and adipokines that impact liver metabolism and inflammation (Lee et al., 2023).

Notably, the liver also impacts the functions of adipose tissue. For example, hepatic peroxisomal β-oxidation has recently been identified as a key regulator of adipocyte browning in diet-induced obesity, through the accumulation of long-chain fatty acids (Lu et al., 2024). The liver plays a critical role in the inter-organ metabolic response to β3-adrenergic stimulation, which leads to triglyceride lipolysis in white adipocytes and thermogenesis activation in brown adipocytes. Activation of β3-adrenergic signaling induces major changes in hepatic gene expression (Fougerat et al., 2022; Simcox et al., 2017). Simcox *et al*. demonstrated that HNF4α activation induces production of liver-derived acylcarnitines that act as alternative fuel for BAT thermogenesis during cold exposure (Simcox et al., 2017). Moreover, β3-adrenergic receptor stimulation induces PPARα- dependent responses in the liver. Hepatocyte PPARα influences insulin secretion and BAT activation induced by β3-adrenergic activation (Fougerat et al., 2022). The interplay between adipose tissue and the liver plays a crucial role in MASLD initiation and progression.

MASLD is a major public health concern; however, research tools are lacking. Preclinical models are essential for not only investigate the disease mechanisms but also for testing potential therapies and identifying novel biomarkers. Existing preclinical mouse models of MASLD are generated using dietary challenges or genetic manipulations, and only partially recapitulate the human disease, exhibiting minimal or no fibrosis, and/or lacking obesity or even exhibiting weight loss (Gallage et al., 2022; Parlati et al., 2021; Smati et al., 2022; Vacca et al., 2024).

Among factors contributing to MASLD development and progression, environmental temperature has emerged as a critical variable in the modeling of human disease. Several studies have highlighted that housing temperature influences mouse physiology (Seeley and MacDougald, 2021). Most rodent studies are conducted at environmental temperatures that correspond to the thermoneutral zone of dressed adult humans, but that are below thermoneutrality for mice. This results in chronic cold stress, which induces catecholamine release by the sympathetic nervous system, leading to brown adipose tissue (BAT) activation and non-shivering thermogenesis to increase heat production and maintain a constant core body temperature. Under these conditions, mice exhibit increases of heart rate, blood pressure, and overall energy expenditure (Fischer et al., 2018; Maloney et al., 2014). Moreover, standard housing suppresses immune responses (Giles et al., 2016; Kokolus et al., 2013). On the other hand, thermoneutral housing accelerates metabolic inflammation in white adipose tissue and in the vasculature during obesity, promoting atherosclerosis progression in mice (Giles et al., 2016; Tian et al., 2016). Housing temperature can impact various diseases and treatment responses (Ganeshan and Chawla, 2017), including tumor growth and responses to immune and cytotoxic therapies (Bucsek et al., 2017; Eng et al., 2015).

As housing temperature has major influences on metabolism and inflammation—which are both involved in MASLD—thermoneutral housing may improve the modeling of preclinical murine models and translatability to the human disease. Few prior studies have investigated how thermoneutral housing influences MASLD development and progression, and the results are controversial. In one study, male mice were housed at thermoneutrality and fed a high-fat diet for 24 weeks, which resulted in accelerated MASLD with enhanced hepatic steatosis, and increased expression of genes involved in hepatic inflammation and fibrosis, compared to standard temperature-housed mice. That study also found that thermoneutral housing enabled obesity and MASLD experimental modeling in female mice, which are typically resistant to obesogenic diet-induced disease development with standard-temperature housing (Giles et al., 2017). Similarly, another study demonstrated that thermoneutral housing of mice exacerbated liver inflammation induced by a diet rich in fat, carbohydrates, and cholesterol (Oates et al., 2023). Moreover, the combination of thermoneutral housing with a high-fat high-fructose diet for mice has been shown to recapitulate many of the histological and genomic characteristics of advanced human MASLD (Morrow et al., 2022). In contrast, another study reported that thermoneutral housing coupled with western diet (WD) feeding did not aggravate diet-induced hepatic inflammation and fibrosis in male or female mice (Nunes et al., 2023). Horakova et al. suggested that the effect of housing temperature may depend on mouse genetic background (Horakova et al., 2023). Overall, the effect of thermoneutrality on diet-induced MASLD remains unclear due to the use of different experimental conditions, including mouse strains, and diet composition and duration.

In the present study, we investigated the impact of thermoneutral housing on a WD-induced mouse model of MASLD (Smati et al., 2022; Vacca et al., 2024). We characterized the effects of environmental temperature on whole-body homeostasis, liver phenotype, and hepatic gene expression. Furthermore, we used MASLD model mice housed at thermoneutrality to investigate the metabolic cross-talk between adipose tissues and the liver during lipolysis induced by β3-adrenergic stimulation.

## RESULTS

### Thermoneutral housing promotes WD-induced obesity, diabetes, and steatosis

To investigate the effect of environmental temperature on diet-induced MASLD, male mice were housed at 22°C (standard temperature, RT) or 30°C (thermoneutral housing, TN), and fed a chow (CD) or western diet (WD) for 13 weeks (Figure 1A). After 5 weeks, WD-fed mice were significantly heavier than CD-fed mice. After 9 weeks, TN-housed WD-fed mice were significantly heavier than RT-housed WD-fed mice (Figure 1B). Housing temperature significantly impacted WD-induced obesity, as confirmed by final total body weight and relative white adipose tissue weight (Figure 1C and 1D). Compared to CD-fed mice, WD-fed mice exhibited significantly lower oral glucose tolerance. Compared to RT-housed WD-fed mice, the TN-housed WF-fed mice showed significantly lower oral glucose tolerance, only at −30 and +30 minutes (Figure 1E and 1F). Fasted insulinemia did not significantly differ between groups (Figure 1G). This resulted in a significantly increased HOMA-IR among TN-housed WD-fed mice only, compared to their CD-fed controls (Figure 1H).

**Figure 1.**
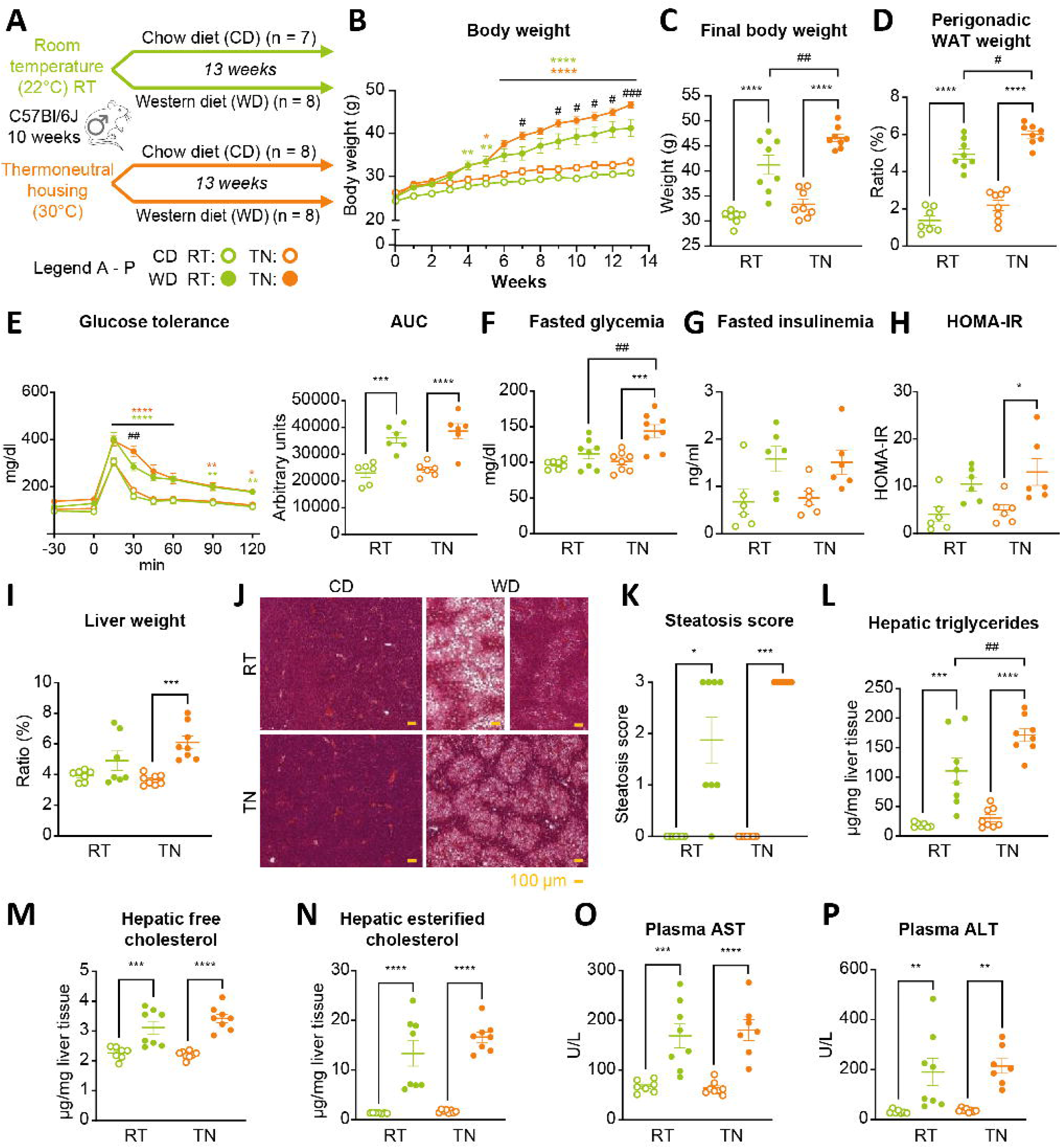
Thermoneutral housing promotes WD-induced obesity, diabetes, and steatosis. (A) WT male C57Bl/6J mice, aged 10 weeks, were housed at room temperature (RT; 22°C) or thermoneutral temperature (TN; 30°C) and fed a chow diet (CD) or western diet (WD) for 13 weeks (*n* = 7–8/group). (B) Body weight was determined weekly during the experiment. (C) Final weight after 13 weeks of diet. (D) Ratio of perigonadal (PG) white adipose tissue (WAT) weight to body weight at the end of the experiment. (E) Oral glucose tolerance test (OGTT) assessed after 10 weeks of diet (*n* = 6/group), and area under the curve (AUC) representing the OGTT results. (F, G) Fasted glycemia (F) and insulinemia (G) measured after 6 hours of fasting. (H) Homeostatic model assessment of insulin resistance (HOMA-IR). (I) Ratio of liver weight to body weight. (J) Representative histological sections of liver, stained with H&E, from each group at 10×. Scale bar, 100 µm. (K) Liver steatosis estimated on histological liver sections. Scoring: parenchymal involvement by steatosis <5%, 0; 5–33%, 1; 33–66%, 2; >66%, 3 (*n* = 7–8/group). (L, M, N) Hepatic neutral lipids (triglycerides, free cholesterol, and esterified cholesterol) extracted from livers, and analyzed by gas-liquid chromatography. (O, P) Plasma alanine aminotransferase (ALT) and aspartate aminotransferase (AST) activity. Data are presented as the mean ± SEM for *n* = 7–8/group. *diet effect; #temperature effect between WD groups; * or #*p* < 0.05; ** or ##*p* < 0.01; *** or ###*p* < 0.001; **** or ####*p* < 0.0001. Differential effects were analyzed by analysis of variance (one or two-way ANOVA) with post-hoc Šídák’s test. Histological scores (K) were analyzed using a non-parametric test (Kruskall- Wallis).

We next investigated the hepatic phenotype. Liver weight was significantly higher in TN-housed WD-fed animals, but did not differ between the groups housed at RT (Figure 1I). Histological H&E staining revealed that WD feeding induced hepatic steatosis, regardless of housing temperature. However, responses to the WD showed more heterogeneity at RT (Figure 1J). The steatosis scores confirmed significant lipid accumulation upon WD at both RT and TN, and the more heterogenous response to WD at RT compared to at TN (Figure 1K). These findings were further supported by the quantification of hepatic triglycerides (Figure 1L). Additionally, both hepatic free cholesterol and esterified cholesterol were higher in the WD group compared to the CD group, with no significant influence of housing temperature (Figure 1M and 1N). Similarly, significantly greater liver damage was observed in WD-fed compared to CD-fed mice, regardless of housing temperature (Figure 1O and 1P).

Overall, these results suggested that TN housing potentiated WD-induced obesity, glucose intolerance, and hepatic steatosis development. However, the effect of TN on these parameters was much less pronounced than that of WD.

### Thermoneutral housing fosters WD-induced changes in hepatic gene expression

We next performed microarray analysis of liver gene expression to identify biological processes sensitive to housing temperature under CD or WD feeding (Table S1). Principal component analysis (PCA) of the whole-liver transcriptome revealed clear separation between the CD-fed and WD-fed groups along the first principal component, accounting for 54% of the variance (Figure 2A). Interestingly, compared to the TN-housed WD-fed animals, the RT-housed WD-fed animals exhibited less clear clustering, with four RT-housed WD-fed individuals clustering close to the CD-fed animals. Our results showed overlap between CD-fed mice at RT and TN, and the same phenomenon was observed among WD-fed mice. Volcano plots of the effect of WD showed that the genomic response to WD was stronger at TN than RT (Figure 2B). Indeed, 705 hepatic genes exhibited significantly modulated expression between WD-fed and CD-fed animals at TN, while only 323 genes were differentially expressed according to WD at RT (Figure 2C). The Venn diagram of these differentially expressed genes (DEGs) showed that among the genes affected by WD at RT, the vast majority were also affected by WD at TN (Figure 2D). Hierarchical clustering using the 739 DEGs confirmed the marked discrimination between WD-fed and CD-fed mice (Figure 2E). Also in accordance with the PCA results, four RT-housed WD-fed individuals were closer to the CD-fed mice than to the other WD-fed mice.

**Figure 2.**
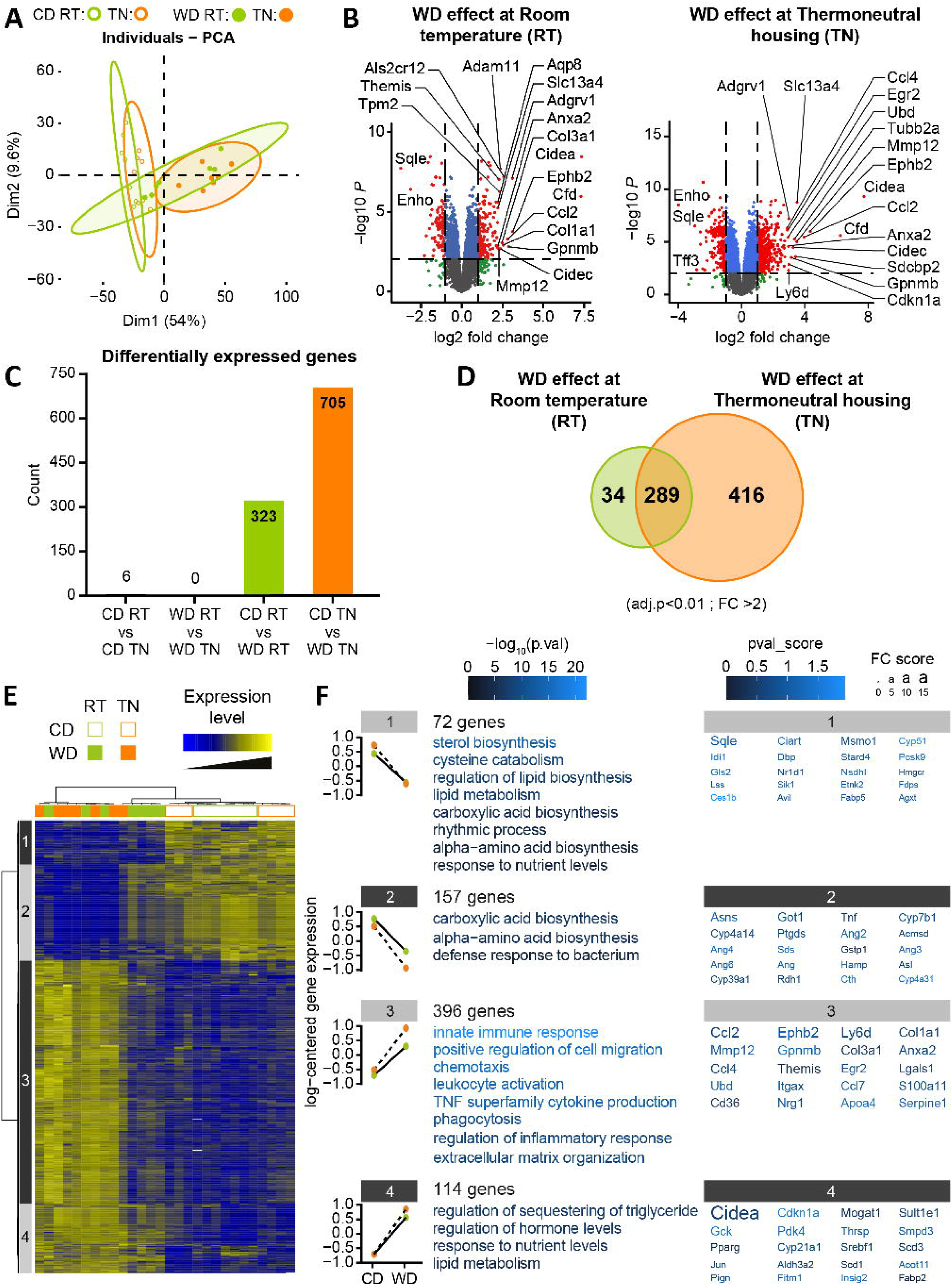
Thermoneutral housing fosters WD-induced changes in hepatic gene expression. (A) Principal component analysis (PCA) score plots of whole-liver transcriptome datasets (*n* = 7– 8/group). Each dot represents an observation (animal), projected onto the first (horizontal axis) and second (vertical axis) PCA variables. (B) Volcano plot shows effects of western diet (WD) on gene expression under room temperature (RT; left panel) or thermoneutral temperature (TN; right panel). Each gene expression level is shown in terms of the –log10 p-value, for comparisons between the WD group and the chow diet group at each temperature. The –log10 *p* values are plotted as a function of the associated log2-fold change, or formally, log2(WD)-log2(Chow). Red indicates *p* < 0.01 and log2(fold change) > 2. Blue indicates *p* < 0.01 and log2 (fold change) < 2. Green indicates *p* > 0.01 and log2(fold change) > 2. Grey indicates *p* > 0.01 and log2(fold change) < 2. Gene names are highlighted for the 20 most highly regulated genes, according to a score based on the adjusted *p* value × logFC. (C) Histogram represents the number of genes significantly regulated for each comparison. (D) Venn diagram represents the number of genes significantly regulated by WD (*p* < 0.001, FC > 2) for each housing temperature (RT or TN). (E) Heatmap represents data from microarray experiments. Significantly differentially expressed genes (adjusted *p* values < 0.01 and fold change > 2) were selected, which corresponded to 739 probes. The color gradient indicates the scaled values of gene expression. Hierarchical clustering identified four gene clusters (indicated on the left). (E) Mean expression profiles for the four gene clusters. Graphs represent the means of the scaled gene expression values. The most significantly enriched biological processes, identified using the Metascape gene ontology algorithm, are shown at the right of each profile. Briefly, hypergeometric tests were performed for each category in each cluster. (F) The top 20 genes in each cluster that showed the largest differences in expression. The color of each character string is related to the *p* value score. The size of each character string is related to the fold change score for all the comparisons made for each gene.

Next, gene clustering revealed four gene clusters along the vertical axis of the heatmap. In the first cluster, 72 genes exhibited lower mRNA expression among WD-fed compared to CD-fed mice, regardless of housing temperature. This cluster was most significantly associated with the biological processes of “sterol biosynthesis” (*p* = 10^−15^) and “regulation of lipid biosynthesis” (*p* = 10^−7^). The most significantly affected genes in cluster 1 included genes involved in cholesterol biosynthesis, such as *Hmgcr*, *Sqle*, *Lss*, and *Msmo1*. The 114 genes in cluster 4 were also upregulated by the WD, regardless of housing temperature. Cluster 4 genes were associated with “sequestering of triglycerides” (*p* = 10^−6^) and “lipid metabolism” (*p* = 10^−6^), and included *Cidea* and *Fitm1*, which are involved in triglyceride storage by promoting lipid droplet formation, and *Thrsp*, which helps regulates the triglyceride biosynthetic process.

On the other hand, the 157 genes in cluster 2 exhibited lower mRNA expression among WD-fed compared to CD-fed mice, which and this different was greater at TN than at RT. Interestingly, among the four RT-housed WD-fed mice that clustered close to the CD-fed animals, the mRNA expression levels of cluster 2 genes were similar to those in the CD-fed mice. Cluster 2 genes were most significantly associated with the biological process of “carboxylic acid biosynthesis” (*p* = 10^−8^). Finally, cluster 3 included 396 genes that exhibited increased expression in WD-fed compared to CD-fed mice, and this difference was temperature-dependent. Notably, as seen for cluster 2, the WD-induced increase in cluster 3. mRNA expression was not observed in the four RT-housed WD-fed mice that clustered with the CD-fed animals. Interestingly, the cluster 3 genes were involved in inflammation and fibrosis with the most significantly affected pathways included “innate immune response” (*p* = 10^−22^), “regulation of inflammatory response” (*p* = 10^−12^), and “extracellular matrix organization” (*p* = 10^−13^). Genes contributing to these pathways included genes encoding for pro-inflammatory cytokines (*Ccl2*, *Ccl4*, and *Ccl7*), for collagens (*Col1a1* and *Col3a1*), and for extra-cellular matrix-degrading metalloproteinases (*Mmp12*) (Figure 2F).

Taken together, these analyses illustrated that WD feeding induced more changes in hepatic gene expression at TN compared to at RT, particularly among genes involved in pro-inflammatory and pro-fibrotic pathways.

### Thermoneutral housing fosters WD-induced hepatic inflammation and fibrosis

Histological analysis was performed to quantify liver inflammation and fibrosis. Blinded assessment of H&E-stained histological sections revealed greater infiltration of inflammatory cells in the liver of WD-fed mice, compared to CD-fed animals (Figure 3A and 3B). Again, mice at RT exhibited a heterogenous response to the WD, which was not observed at TN. However, the inflammation score was only significantly higher in TN-housed WD-fed mice, compared to TN-housed CD-fed mice (Figure 3B). These findings correlated with the increased hepatic expressions of *Tnfα* and *Il1β* (Figure 3C). We next assessed fibrosis by alpha smooth muscle cell actin (α-SMA) staining, which indicated stellate cell activation, as well as Sirius red staining, which marked type I and II collagen fibers (Figure 3D and 3G). Both α-SMA and collagen were significantly more abundant in TN-housed WD-fed mice, compared to TN-housed CD-fed mice, while we observed no significant difference between WD-fed and CD-fed mice at RT (Figure 3E 3H). Notably, the combination of WD and TN was the only experimental condition in which we observed a fibrosis score of 2. These findings correlated with the increased gene expressions of *Acta2*, *Mmp13*, *Col1a1*, and *Col3a1* in TN-housed WD-fed mice (Figure 3F and 3I).

**Figure 3.**
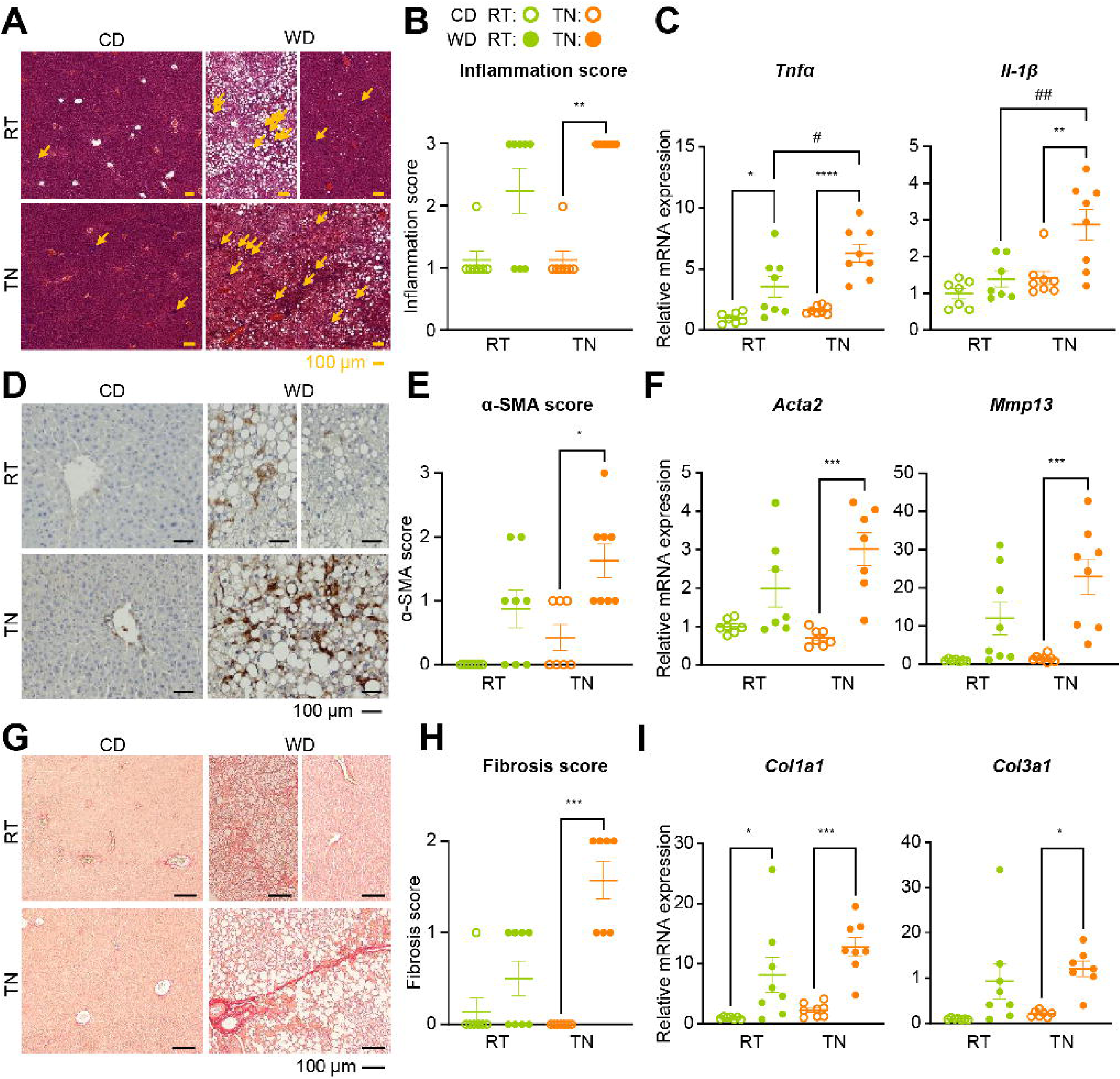
Thermoneutral housing fosters WD-induced hepatic inflammation and fibrosis. (A) Representative histological sections of liver stained with H&E from each group at 10x. Scale bar, 100 µm. (B) Liver inflammation estimated on histological liver sections. H&E staining was used to evaluate inflammation as follows: 0, no focus; 1, more than 2 foci; 2, 2–4 foci; 3, more than 4 foci (*n* = 7– 8/group). (C) Hepatic mRNA expression of inflammatory genes (*Tnfα* and *Il-1β*) measured by RT-qPCR. (D) Representative histological sections of liver from each group, evaluated by immunohistochemical staining with alpha smooth muscle (α-SMA), at 20×. Scale bar, 100 µm. (E) Stellate hepatic cell activation estimated on histological liver sections. Scoring: <3% of periportal area, 0; 3–33%, 1; 34–66%, 2; >66%, 3 (*n* = 7–8/group). (F) Hepatic mRNA expression of genes involved in stellate cell activation (*Acta2* and *Mmp13)* measured by RT-qPCR. (G) Representative histological sections of liver from each group, stained with Sirius red, at 10x. Scale bar, 100 µm. (H) Liver fibrosis estimated on histological liver sections. Sirius red staining was used to evaluate fibrosis as follows: 0, no fibrosis; 1, pericellular and perivenular fibrosis; 2, focal bridging fibrosis (*n* = 7–8/group). (I) Hepatic mRNA expression of fibrosis genes (*Col1a1* and *Col3a1*) measured by RT-qPCR. Data are presented as the mean ± SEM for *n* = 7–8/group. *diet effect; #temperature effect; * or #*p* < 0.05; \*\**p* < 0.01; *** or ###*p* < 0.001; \*\*\*\**p* < 0.0001. Differential effects were analyzed by analysis of variance (one-way ANOVA) with post-hoc Šídák’s test (B, E, H) Histological scores were analyzed using a non-parametric test (Kruskall-Wallis).

Altogether, these results revealed that TN housing aggravated WD-induced inflammation and fibrosis, suggesting an acceleration of MASLD progression.

### Thermoneutral housing reduces the intra-group variability in WD-induced MASLD

Upon finding a more heterogenous response to WD feeding under RT compared to TN conditions, we wondered whether the heterogenous response observed at RT might be due to a cage effect. However, each cage of RT-housed WD-fed mice included two individuals that were “bad responders” and two that were “good responders”, in terms of body weight, liver damage, hepatic steatosis, and fibrosis (Figure S1). To further investigate the TN housing-induced decrease in heterogeneity of response to WD, we first analyzed the hepatic gene variance distribution in our microarray data. Among CD-fed mice, the distribution of gene variances overlapped between RT-and TN-housed mice. On the other hand, among WD-fed mice, the RT-housed mice exhibited a shift of the variance distribution to the right side of the plot, indicating higher intra-group variance in this group compared to among TN-housed mice (Figure S2A). This was confirmed by a decreased mean intra-group variance for the non-homoscedastic genes (i.e., genes showing a significant difference in the intra-group variance between the experimental groups) in TN-housed WD-fed mice compared to RT-housed WD-fed mice, while no significant difference was observed between the groups of CD-fed mice (Figure S2B). Interestingly, most of the genes showing significantly higher intra-group variability in RT-housed WD-fed animals were related to inflammation and fibrosis (Figure S3). Finally, we confirmed that in terms of body weight, liver damage, and hepatic steatosis, the intra-group standard error of mean (SEM) was significantly lower among TN-housed WD-fed mice compared to RT-housed WD-fed mice (Table S2). Overall, this analysis demonstrated that compared to RT housing, TN housing significantly reduced the intra-group variability in response to WD feeding.

### WD-induced MASLD is associated with altered hepatic gene expression in response to β3- adrenergic stimulation

During MASLD development, free fatty acids (FFA) released by white adipose tissue (WAT) lipolysis are the major source of fatty acids for hepatic lipid storage (Donnelly et al., 2005; Fabbrini et al., 2008). Moreover, we recently demonstrated that adipocyte ATGL-dependent lipolysis controls liver gene expression in response to fasting or β3-adrenergic stimulation (Fougerat et al., 2022). However, the integrity of this WAT-to-liver axis in MASLD has not been well-investigated. Thus, we took advantage of our nutritional MASLD model to investigate this inter-organ dialogue. We induced adipocyte lipolysis through β3-adrenergic receptor activation using CL316243 (CL) and fasting, in male mice housed at TN and fed the CD or WD (Figure 4A). After CL treatment, both CD- fed and WD-fed mice exhibited significantly increased FFA and glycerol levels (Figure 4B). Both diet groups showed decreased blood glucose levels following CL treatment, but the insulin levels after CL-induced β3-adrenergic signaling activation were higher in WD-fed mice than in CD-fed mice (Figure 4C).

**Figure 4.**
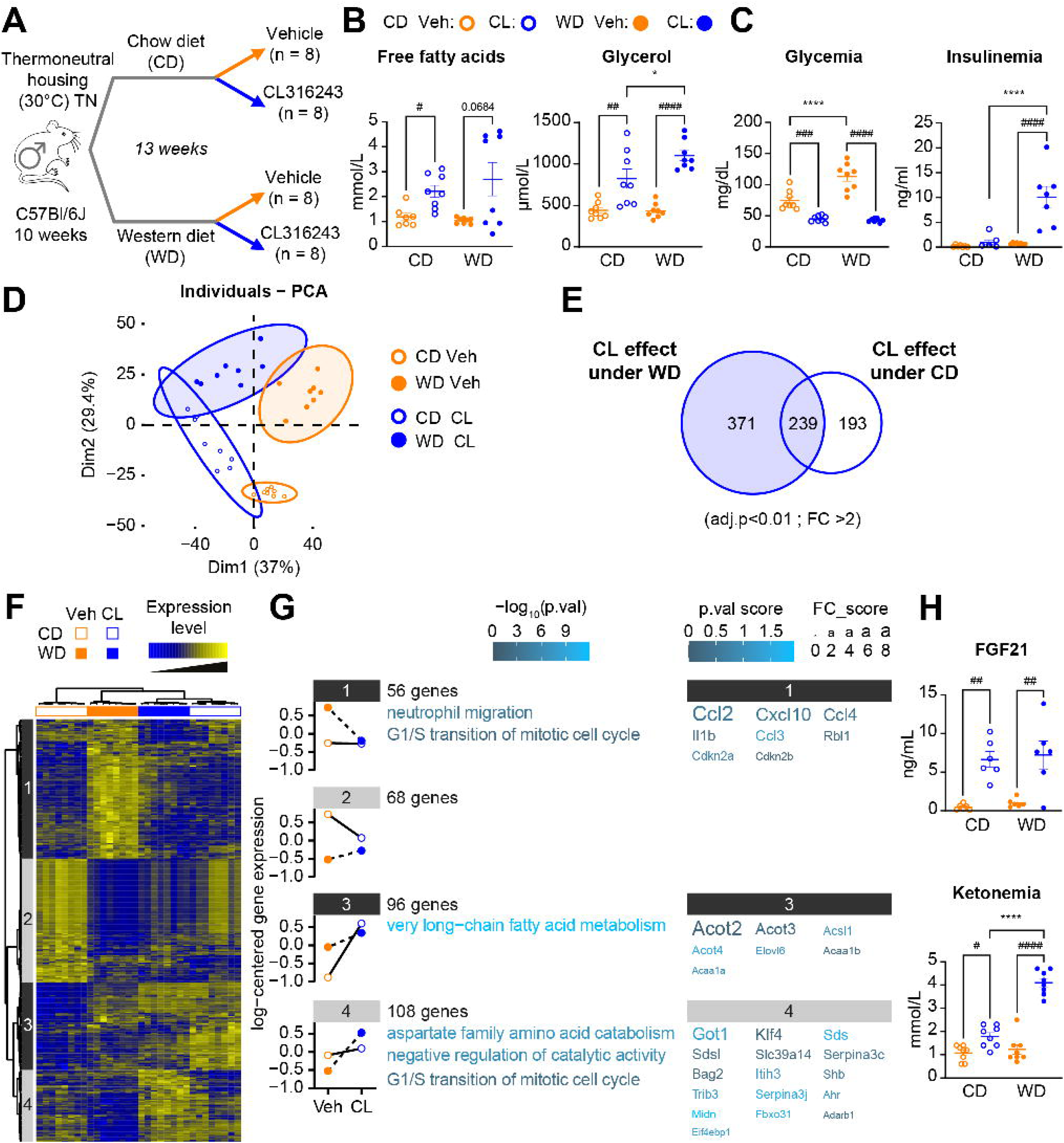
WD-induced MASLD is associated with altered hepatic gene expression in response to β3-adrenergic stimulation. (A) WT male C57Bl/6J mice, aged 10 weeks, housed at thermoneutral temperature (TN; 30°C) for 13 weeks and fed a chow diet (CD) or western diet (WD). At the end of experiment, mice fasted at ZT0, were given CL316243 or vehicle by gavage at ZT10, and were sacrificed at ZT16. (B) Plasma free fatty acid and glycerol levels. (C) Blood glucose and plasma insulin levels at sacrifice. (D) Venn diagram represents the number of genes significantly regulated by CL for each diet (CD or WD). (E) Principal component analysis (PCA) score plots of the whole-liver transcriptome datasets (*n* = 8/group). Each dot represents an observation (animal), projected onto the first (horizontal axis) and second (vertical axis) PCA variables. (F) Heatmap represents data from microarray experiments. Selected genes were significantly differentially expressed, with interaction between the CL effect and the diet (adjusted *p* values < 0.05 and fold change > 1.5), which corresponded to 332 probes. The color gradient indicates the scaled values of gene expression. Hierarchical clustering identified four gene clusters (indicated on the left). (E) Mean expression profiles for the four gene clusters. Graphs represent the means of the scaled gene expression values. The most significantly enriched biological processes, identified using the Metascape gene ontology algorithm, are shown at the right of each profile. Briefly, hypergeometric tests were performed for each category in each cluster. (H) Circulating levels of Fgf21 and ketone bodies (β-hydroxybutyrate). Data are presented as the mean ± SEM for *n* = 8/group. *diet effect; #CL316243 effect; * or #*p* < 0.05; \*\**p* < 0.01; *** or ###*p* < 0.001; **** or ####*p* < 0.0001. Differential effects were analyzed by analysis of variance (one-way ANOVA) with post-hoc Šídák’s test.

We next assessed liver gene expression using microarray analysis, to identify biological processes that were sensitive to CL treatment under CD or WD feeding (Table S3). PCA of the whole hepatic transcriptome revealed clear separation between the vehicle and CL groups along the first principal component, accounting for 37% of the variance. We also observed discrimination between the CD-fed and WD-fed groups along the second principal component, accounting for 29.4% of the variance (Figure 4D). A Venn diagram showed that CL treatment affected a set of genes shared among CD-fed and WD-fed mice, but also diet-specific genes, especially in response to WD (Figure 4E). Therefore, we performed hierarchical clustering using the 332 genes that displayed a significant interaction between CL treatment and diet (Figure 4F). The gene clustering revealed four major genetic groups along the vertical axis of the heatmap (Figure 4G). In cluster 1, 56 genes displayed lower expression in CL-treated vs. vehicle-treated mice, only among WD-fed mice. These genes were notably involved in the biological process of “neutrophil migration” (*p* = 10^−5^), and included genes for chemokines, such as *Ccl2*, *Cxcl10*, and *Ccl4*. Similarly, cluster 4 included 104 genes that showed higher expression in CL-treated than in vehicle-treated mice, only among WD-fed mice. This cluster showed involvement in “aspartate family amino acid catabolism” (*p* = 10^−5^). In contrast, the 68 genes from cluster 2, and the 96 genes from cluster 3, were significantly impacted upon CL treatment in CD-fed mice, but not in WD-fed mice. Interestingly, cluster 3 genes that exhibited higher expression in CL-treated vs. vehicle-treated CD-fed mice were involved in “very long-chain fatty acid metabolism” (*p* = 10^−10^). Genes contributing to this pathway included *Acot1–4*, encoding acyl-coA thioesterases, which have been described as target genes of peroxisome proliferator-activated receptor α (PPARα) upon fasting (Fougerat et al., 2022; Montagner et al., 2016). On the other hand, PPARα-dependent responses, such as ketone body and FGF21 production, were increased in response to CL-induced adipocyte lipolysis among both CD-fed and WD-fed mice (Figure 4H).

Collectively, these results demonstrated that CL-induced β3-adrenergic stimulation induced WAT lipolysis, and PPARα-dependent changes in liver gene expression and metabolism. However, MASLD entails alteration of some of the gene expression changes that occur in response to β3- adrenergic stimulation.

### WD-induced MASLD is associated with defective BAT activation in response to β3-adrenergic stimulation

β3-adrenergic signaling induces triacylglycerol lipolysis in WAT, as well as activates thermogenesis in BAT (Blondin et al., 2017; Haemmerle et al., 2006; Labbé et al., 2015; Li et al., 2014). Again, we took advantage of our TN-housing WD-induced MASLD preclinical model to investigate the consequences of MASLD on BAT activation, in response to high-level lipolysis induced by combined CL treatment and fasting (Figure 4A). Compared to their vehicle counterparts, CL-treated CD-fed mice exhibited significantly lower BAT weight, while CL-treatment did not significantly impact BAT weight among WD-fed mice (Figure 5A). Microarray analysis of BAT gene expression revealed biological processes that were sensitive to CL treatment under CD or WD feeding (Table S4). PCA of the whole BAT transcriptome showed clear separation between vehicle- and CL-treated mice along the first principal component, accounting for 48.2% of the variance. On the other hand, no significant discrimination was observed between CD-fed and WD-fed mice on any principal component (Figure 5B). However, a Venn diagram showed that CL treatment affected a much higher number of genes in CD-fed mice (2701 DEGs) than in WD-fed mice (1688 DEGs) (Figure 5C). We confirmed that *Elovl3* mRNA expression was significantly decreased in CL-treated WD-fed mice compared to CL-treated CD-fed mice, and that other BAT markers exhibited a lower fold change of induction upon CL treatment among WD-fed mice compared to CD-fed mice (Marcher et al., 2015) (Figure S4).

**Figure 5.**
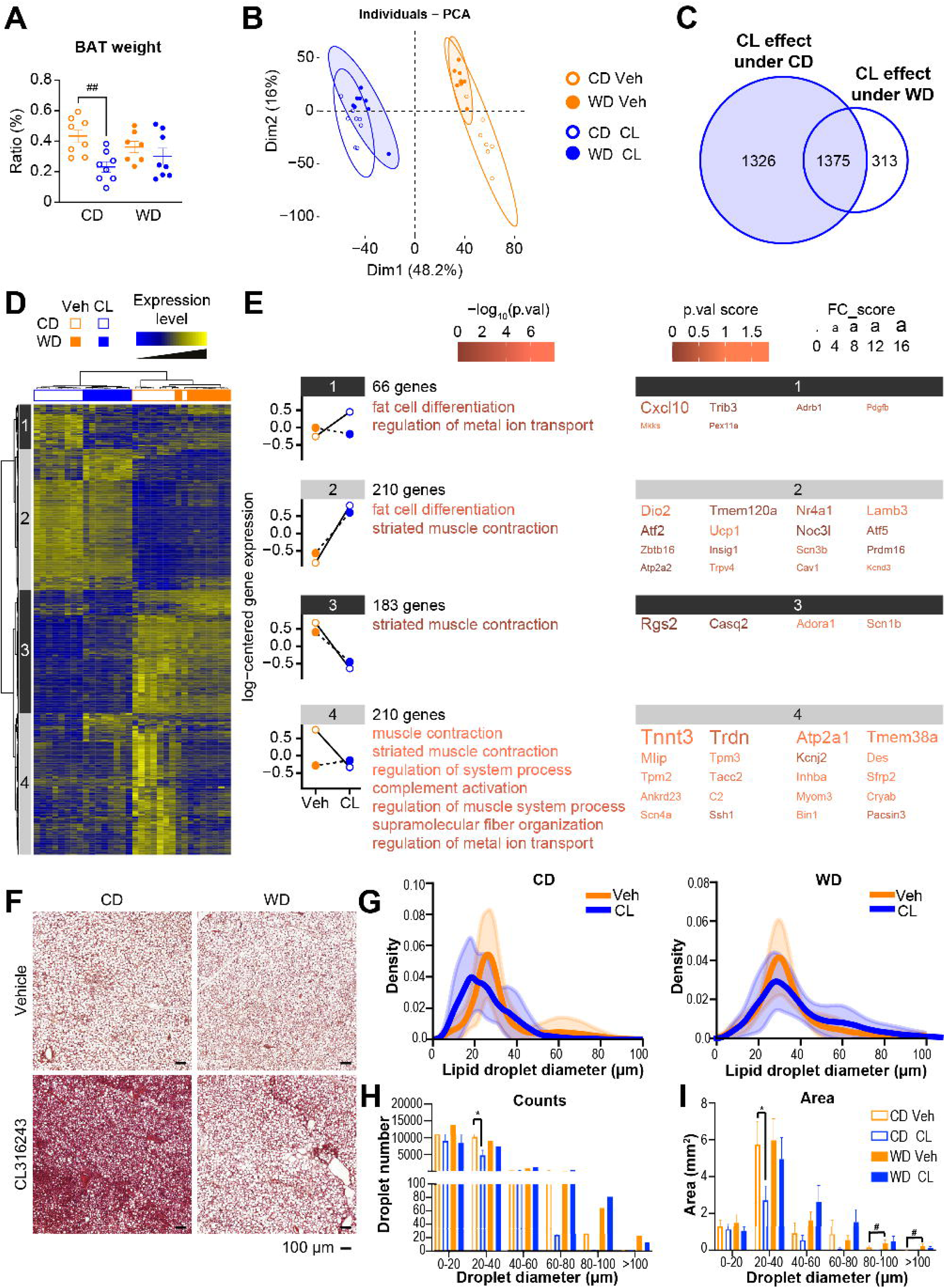
WD-induced MASLD is associated with defective BAT activation in response to β3- adrenergic stimulation. (A) Ratio of brown adipose tissue (BAT) to body weight. (B) Principal component analysis (PCA) score plots of the whole-BAT transcriptome datasets (*n* = 8/group). Each dot represents an observation (animal), projected onto the first (horizontal axis) and second (vertical axis) PCA variables. (C) Venn diagram represents the number of genes significantly regulated by CL for the chow diet (CD) or western diet (WD). (D) Heatmap represents data from microarray experiments. The selected genes were significantly differentially expressed, with interaction between the CL effect and the diet (adjusted *p* values < 0.05 and fold change > 1.5), which corresponded to 669 probes. The color gradient indicates the scaled values of gene expression. Hierarchical clustering identified four gene clusters (indicated on the left). (E) Mean expression profiles for the four gene clusters. Graphs represent the means of the scaled gene expression values. The most significantly enriched biological processes, identified using the Metascape gene ontology algorithm, are shown at the right of each profile. Briefly, hypergeometric tests were performed for each category in each cluster. (F) Representative histological sections of BAT stained with H&E from each group at 10x. Scale bar 100 µm. (G) Lipid droplet distribution according to their size. (H) Lipid droplet counts. (I) Lipid droplet area. Data are presented as the mean ± SEM for *n* = 8/group. *diet effect; #CL316243 effect; * or #*p* < 0.05; \*\**p* < 0.01; *** or ###*p* < 0.001; **** or ####*p* < 0.0001. Differential effects were analyzed by analysis of variance (one-way ANOVA) with post-hoc Šídák’s test. (F) Histological scores were analyzed using a non-parametric test (Kruskall-Wallis).

We performed hierarchical clustering using the 669 genes showing a statistically significant interaction between CL treatment and diet, which confirmed the marked discrimination between CL-treated and vehicle-treated mice, regardless of diet, as previously observed by PCA (Figure 5D). Subsequent gene clustering revealed four major genetic groups along the vertical axis of the heatmap. Interestingly, in all clusters, the effect of CL treatment observed in CD-fed mice was significantly lower (clusters 2 and 3) or completely blunted (clusters 1 and 4) in WD-fed mice. Among the genes that exhibited a decreased response to CL-induced lipolysis in WD-fed mice, some were involved in “fat cell differentiation” (clusters 1 and 2), including genes related to brown adipocyte differentiation and thermogenesis, such as *Dio2*, *Ucp1*, and *Prdm16* (Figure 5E). Histological staining revealed lower numbers of lipid droplets in CL-treated CD-fed mice compared to in vehicle-treated CD-fed mice; however, this CL-dependent effect was not observed in WD-fed mice (Figure 5F). This finding was further confirmed by measurement of lipid droplet diameter, which revealed a shift towards smaller lipid droplets in the BAT of CL-treated CD-fed mice, compared to vehicle-treated CD-fed mice. In contrast, these two distributions overlapped among WD-fed mice (Figure 5G). Comparisons of the number of lipid droplets aggregated according to size, and of the area occupied by the droplets, further illustrated that CL-treatment significantly decreased the number lipid droplets and the surface occupied by small lipid droplets (20–40 μm) among CD-fed mice, but not WD-fed mice (Figure 5H and 5I). Overall, BAT activation in response to CL-induced β3-adrenergic stimulation was blunted in this mouse model of obesity and MASLD.

MASLD has become the most prevalent chronic liver disease worldwide, with a 30% prevalence in the adult population (Riazi et al., 2022; Rinella et al., 2023; Younossi et al., 2019). Since diet-induced obesity is a major risk factors driving MASLD development, dietary-based preclinical models are used to investigate development of the human pathology. The so-called “western diets”, which are high in fat and carbohydrates, are generally accepted as relevant models (Gallage et al., 2022; Vacca et al., 2024). However, mounting evidence shows that mouse models housed at a suboptimal, col-stress-inducing, ambient temperature (22°C) do not recapitulate human metabolic pathologies (Feldmann et al., 2009; Ganeshan and Chawla, 2017; Škop et al., 2020; Vialard and Olivier, 2020; X et al., 2003). Combining diet-induced obesity with thermoneutral housing has emerged as a promising method of refining the mouse model of MASLD. However, previous studies of C57Bl6 mice have induced MASLD using a high-fat diet combined with TN housing, and have failed to achieve robust hepatic fibrosis, a key feature of human MASLD (Giles et al., 2017; Horakova et al., 2023). In the present study, we combined TN housing with a WD enriched in fat, sucrose, and cholesterol, and observed consistent aggravation of diet-induced metabolic perturbations. WD-fed TN-housed mice presented with higher body weight and adiposity, and decreased glucose tolerance, compared to WD-fed RT-housed mice. Interestingly, WD-fed TN-housed mice had higher hepatic steatosis, compared to their RT-housed counterparts. These mice even developed liver inflammation and fibrosis—confirmed by both gene expression and histological studies—thereby confirming that our MASLD model mirrored pathological and molecular events consistent with human pathogenesis. Our results are consistent with those of Morrow (Morrow et al., 2022), which also demonstrated that TN housing significantly promoted histological and molecular MASLD features induced by a high-fat high-fructose diet. However, several other authors have reported no significant differences in steatosis or fibrosis development between WD-fed mice housed at TN or RT (Nunes et al., 2023; Oates et al., 2023). This discrepancy is probably not due to differences in the duration of the dietary interventions, since all above-cited studies were performed for at least 16 weeks, like our study. It is more likely that specific nutrients within the different diets might interact with the housing temperature to promote inflammation and fibrosis in the liver. Indeed, a recent study highlighted the critical role of diet composition in modeling the different steps of MASLD progression at thermoneutrality (Low et al., 2024).

Interestingly, our study revealed that TN housing decreased the intra-group variability in response to WD feeding, compared to housing at RT. This difference was observed for most investigated physiological parameters, including body weight, plasmatic enzymes, liver steatosis and inflammation, and hepatic gene expression. Consistent with our phenotypic observations, the genes exhibiting the most heterogenous responses at RT were involved in inflammation and fibrosis development. Our findings are in agreement with previous reports that TN promotes inflammation in the vasculature (Tian et al., 2016), adipose tissue (Giles et al., 2016), and liver (Giles et al., 2017; Morrow et al., 2022). Moreover, our results suggest that compared to RT, TN induces a more homogenous response to the WD, and an accelerated response to the WD, at least in the liver. Importantly, by reducing the interindividual variability in response to dietary challenges, TN housing might enable significant reduction of the required duration of the dietary intervention and/or of the number of animals necessary to model MASLD. Notably, the number of mice per cage has been shown to influence thermoregulation, through regulation of *Ucp1* expression in the BAT (Škop et al., 2021).

In addition to alterations of intrinsic liver metabolism, dysfunctional WAT strongly contributes to MASLD initiation and progression. WAT is the major site of lipid storage, and releases lipids into the circulation to fuel other organs (especially the liver) in case of energy need, thereby regulating systemic energy homeostasis (Rosen and Spiegelman, 2006). In obesity, WAT becomes dysfunctional, leading to increased basal lipolysis, dysregulated adipokine secretion, and release of pro-inflammatory molecules. In the liver, these mechanisms yield increased hepatic lipid content and inflammation, which further promote insulin resistance (Lee et al., 2023). In the present study, we used our TN-housed MASLD pre-clinical model to investigate the metabolic cross-talk between adipose tissues and the liver. We first observed that WD-fed mice did not exhibit impaired induction of adipose lipolysis by acute activation of β3-adrenergic signaling. Accordingly, the CL-mediated increase of plasma insulin secretion, which depends on adipose lipolysis (Heine et al., 2018), was also not altered, and was even higher in mice with WD-induced MASLD compared to healthy mice. We recently demonstrated that ATGL-dependent lipolysis is a regulator of hepatic functions via control of liver gene expression upon β3-adrenergic stimulation (Fougerat et al., 2022).

Activation of β3-adrenergic signaling induces major changes in hepatic gene expression (Fougerat et al., 2022; Simcox et al., 2017). Interestingly, in our present study, gene expression analysis revealed a set of hepatic genes regulated upon acute lipolysis stimulation, mostly in CD-fed mice and less in WD-fed mice, suggesting that these genes were less sensitive to CL in WD-induced MASLD. Some of these genes are known targets of PPARα, a nuclear receptor highly expressed in hepatocytes, which regulates the expression of genes controlling hepatic fatty acid catabolism during fasting (Kersten et al., 1999; Montagner et al., 2016; Régnier et al., 2018). We recently identified hepatocyte PPARα as a key player of the adipose-to-liver dialog that is required for hepatic gene expression, ketogenesis, and FGF21 production in response to WAT lipolysis (Fougerat et al., 2022). However, our current results showed that ketogenesis and FGF21 production—two processes under the transcriptional control of PPARα in hepatocytes (Badman et al., 2007; Inagaki et al., 2007; Montagner et al., 2016; Régnier et al., 2018)—were similarly induced after β3-adrenergic receptor stimulation in CD-fed and WD-fed mice, indicating that these two hepatocyte PPARα-dependent hepatic functions remained functional in WD-induced MASLD. These findings suggest that the WAT lipolysis-dependent activity of hepatic PPARα is intact in MASLD. However, we cannot exclude that other lipolysis stimulated-hepatic pathways—such as HNF4α-dependent gene expression—may be altered in WD-induced MASLD (Simcox et al., 2017).

Finally, our results revealed that BAT activation in response to β3-adrenergic signaling stimulation was reduced in WD-induced MASLD, suggesting altered cross-talk between the liver and the BAT. This finding is consistent with a previous report that diet-induced obesity mice show reduced uptake of thermogenesis substrates (especially triglyceride-rich lipoproteins) by the BAT during β3-adrenergic receptor stimulation (Heine et al., 2018). To ensure efficient thermogenesis, brown adipocytes also use other energy-rich substrates, including glucose (Hankir and Klingenspor, 2018; Orava et al., 2011; Yu et al., 2002), FFAs released by white adipocytes (Schreiber et al., 2017; Shin et al., 2017), and acylcarnitines (Simcox et al., 2017). In our present study, we did not analyze substrate uptake into BAT; however, we did not detect differences in the triglyceride, glucose, or FFA plasma levels in WD-fed mice compared to CD-fed mice.

In response to acute adipocyte lipolysis, the transcription factors PPARα and HNF4α not only control hepatic gene expression but also influence BAT activation (Fougerat et al., 2022; Simcox et al., 2017). Our present findings indicated that WD feeding did not drastically influence PPARα-dependent signaling, suggesting that reduced BAT activation in WD-induced MASLD does not depend on hepatocyte PPARα activity. However, we cannot exclude the possibility that hepatocyte HNF4α plays a role in the effect of WD on BAT activation.

As already suggested, we assumed that β3-adrenergic signaling activation promotes insulin secretion from pancreatic β cells (Cypess et al., 2015; Heine et al., 2018). This response to CL depends on adipose lipolysis, and is required for BAT activation (Heine et al., 2018). Upon activation of β3-adrenergic signaling, we found that plasma insulin levels were not altered in WD-fed mice, and were even higher compared to CD-fed mice. However, despite a high increase of plasma insulin levels, BAT activation was reduced in mice with WD-induced MASLD, suggesting defective BAT insulin signaling. Accordingly, mice with diet-induced obesity exhibit insulin resistance in BAT, and reduced BAT glucose uptake in response to β3-adrenergic signaling activation (Roberts-Toler et al., 2015). Moreover, in a study using mice with an inducible brown adipocyte-specific insulin receptor deletion, Heine *et al*. showed that insulin signaling in BAT is essential for energy-rich substrate uptake and thermogenesis by BAT in response to β3-adrenergic stimulation (Heine et al., 2018). Although we cannot exclude other hepatic or extra-hepatic mechanisms, our present findings suggest that insulin resistance in BAT likely contributes to the reduced BAT activation in WD-induced MASLD.

In conclusion, our present findings demonstrated that the combination of thermoneutral housing and WD feeding produces an accelerated mouse model of MASLD, which could facilitate the elucidation of the molecular mechanisms underlying the steatosis-to-steatohepatitis transition, as well as the testing of therapeutic approaches. We further used this model to show that MASLD is associated with an altered hepatic response to white adipose tissue lipolysis, including changes in hepatic gene expression, enhanced insulin levels, and reduced BAT activation in response to β3- adrenergic stimulation. Altogether, these results provide a useful resource for the modeling of MASLD and metabolic dialogue with adipose tissues.

## Limitations of the study

A first limitation of our study is that we did not evaluate females. We demonstrated that thermoneutral housing combined with WD feeding promoted MASLD in male mice; however, MASLD is a sexually dimorphic disease. Additionally, we used our MASLD preclinical mouse model to assess the metabolic response to β3-adrenergic activation between the liver and the adipose tissues. While we provided evidence that BAT response to β3-adrenergic stimulation was reduced in WD-induced MASLD, we did not identify the mechanisms that drive this effect.

## STAR Methods

### Key resources table

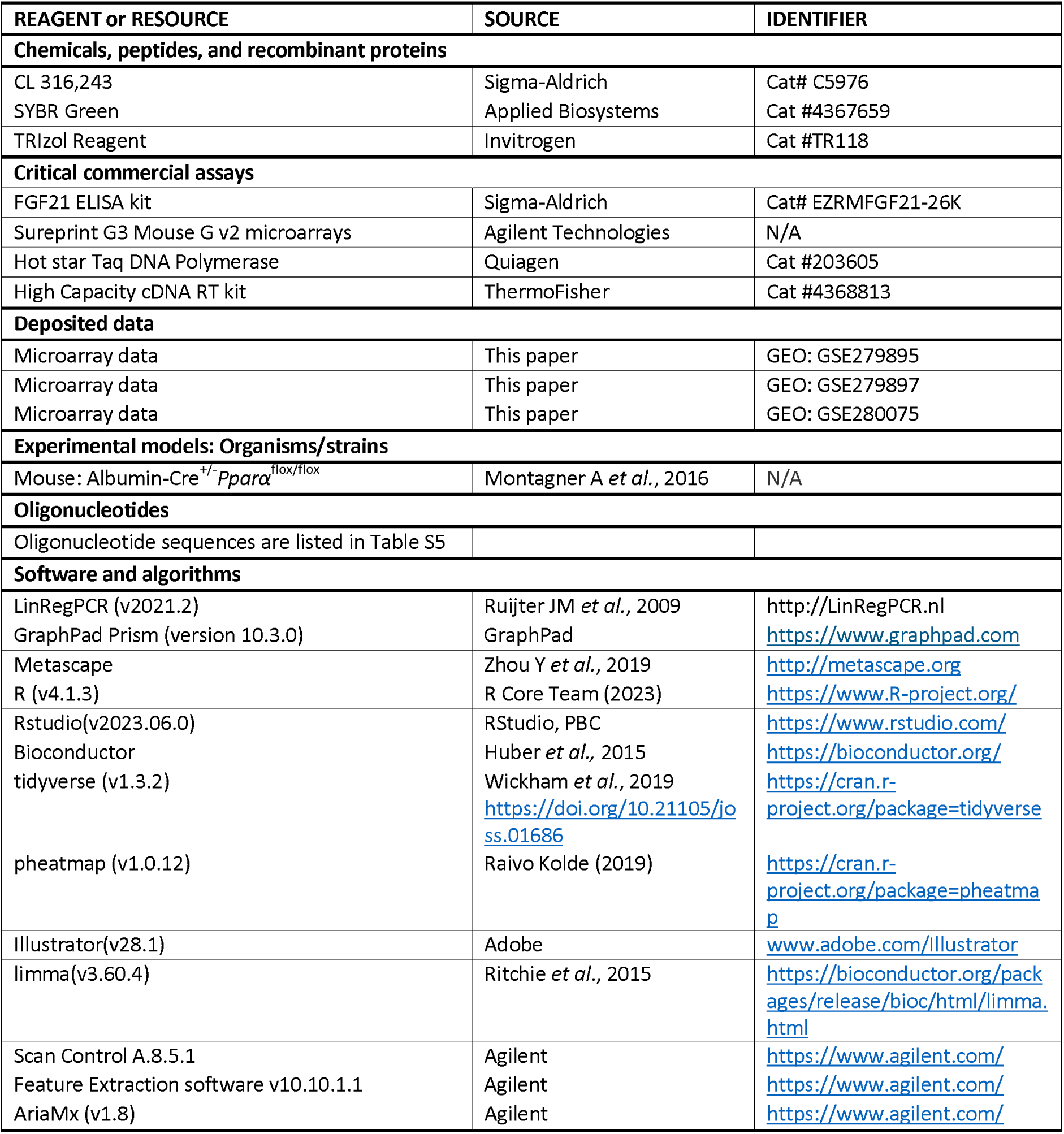

### Resource availability Lead contact

Further information and requests for resources and reagents should be directed to and will be fulfilled by the lead contact, Nicolas Loiseau.

### Material availability

This study did not generate new reagents or Mouse lines for this study.

### Method details Animals

All experiments were carried out in accordance with the European Guidelines for the Care and Use of Animals for Research Purposes and approved by an independent ethics committee (CEEA-86 Toxcométhique) under the authorization number 17430-2018110611093660 v3. The animals were treated humanely with due consideration to the alleviation of distress and discomfort. A total of 64 C57BL/6J male mice (8-9-week-old) were purchased from Charles Rivers Laboratories (L’Arbresle, France). All mice were housed on a 12 h light (ZT0-ZT12)/12 h dark (ZT12-ZT24) cycle in a ventilated cabinet at the specific temperature, either a standard temperature (RT = 21-23°C) or a thermoneutral temperature (TN = 28-30°C) throughout the experiment.

Mice were allowed two weeks of acclimatization with free access to water and food (standard rodent diet (safe A04 U8220G10R) from SAFE (Augy, France)). Then, the mice were randomly divided into four groups of 8 mice each. Two groups (n=16, 4 cages of 4 mice) were fed a chow diet (CD, A04, Safe Diets) and the other two groups (n=16, 4 cages of 4 mice) were fed a Western diet (WD, TD.88137, Envigo) for 13 weeks. Half of CD (n=8, 2 cages of 4 mice) and WD (n=8, 2 cages of 4 mice) groups were housed at TN while the other half were housed at RT for 13 weeks. Experimental groups were designed as follows: CD RT, WD RT, CD TN, WD TN. WD contains 42% calories from fat, 42.7% calories from carbohydrates (mainly sucrose) and 15.2% calories from proteins and 0.2% cholesterol; or a CD contains 8.4% calories from fat, 19.3% calories from carbohydrates and 72.4% calories from proteins. Body weight, food and water intake were measured weekly. At the end of the experiment, mice were sacrificed at ZT16 to collect plasma and tissue samples as described below.

### Oral glucose tolerance test and plasma insulin measurement

All experiments were performed using conscious mice. After 10 weeks of feeding, six mice per group were randomly chosen, then fasted for 6h before receiving an oral glucose load (2 g/kg body weight). Blood glucose was measured at the tail vein using an Accu-Check® Performa glucometer (Roche Diabetes Care France, Mylan, France) at 30 min before and 0, 15, 30, 60, 90 and 120 min after the glucose load. At 30 min before and 15 min after glucose injection, 20 µl blood from the tip of the tail vein was sampled for measurement of plasma insulin concentration using HTRF serum kits (Cisbio, Codolet, France). Briefly, 5µl serum was incubated overnight at 4°C with the two corresponding monoclonal antibodies. When dyes are in close proximity, donor excitation via a light source triggers a fluorescence resonance energy transfer (FRET). The fluorescence energy transfer was measured using a Tecan Infinite 500 plate reader (Tecan, Lyon, France). The results were analyzed against a standard curve fit with the four-parameter logistic (4 PL) model, following the instructions for the kit (GraphPad Prism, USA).

### β3-adrenergic receptor activation

Sixteen 23-week-old mice were fasted at ZT0 and given CL316243 (3 mg/kg body weight; Sigma Aldrich) or vehicle (0.5% carboxymethylcellulose in sterilized water) by gavage at ZT10 and sacrificed at ZT16 (n = 8 per group).

### Blood and tissue sampling

Prior to sacrifice, the submandibular vein was lanced and blood was collected into lithium heparin-coated tubes (BD Microtainer®, BD, Dutscher, Brumath, France). Then, mice were killed by cervical dislocation. Plasma was isolated by centrifugation (1500 × g, 15 min, 4°C) and stored at -80°C until biochemical analysis. Tissue samples (liver, subcutaneous and perigonadic white adipose tissue (WAT), brown adipose tissue (BAT), caecum) were collected, weighed, dissected (when necessary) and prepared for histology analysis or snap-frozen in liquid nitrogen and stored at −80°C until further analyses.

### Biochemical Plasma analysis

Plasma samples were assayed for aspartate transaminase (AST), alanine transaminase (ALT), alkaline phosphatase (ALP), free fatty acids (FFAs), triglycerides (TGs), total cholesterol, low-density lipoprotein (LDL-Cholesterol) and high-density lipoprotein (HDL-Cholesterol). All biochemical analyses were performed with a ABX Pentra 400 biochemical analyzer (Anexplo facility, Toulouse, France). Blood glucose levels were measured from mandibular vein using AccuCheck Performa glucometer strips (Roche Diagnostics), while β-hydroxybutyrate content was measured using Optium β-ketone test strips bearing Optium Xceed sensors (Abbott Laboratories, Abbott Park, IL, USA).

### Liver neutral lipid analysis

Hepatic lipids contents were extracted as previously described (BLIGH and DYER, 1959). Briefly, tissue samples were homogenized in Lysing Matrix D tubes with 1ml methanol/5mM EGTA (ethylene glycol-bis(β-aminoethyl ether)-N,N,N’,N’-tetraacetic acid) (2:1, v/v) in a FastPrep machine (MP Biochemicals). Lipids (corresponding to an equivalent of 2 mg of tissue) were extracted in chloroform/methanol/water (2.5:2.5:2, v/v/v), in the presence of the following internal standards: glyceryl trinonadecanoate, stigmasterol, and cholesteryl heptadecanoate (Sigma-Aldrich, Saint-Quentin-Fallavier, France). Total lipids were suspended in 160 µl ethyl acetate, and the triglycerides, free cholesterol, and cholesterol ester were analyzed with gas- chromatography on a Focus Thermo Electron system using a Zebron-1 Phenomenex fused-silica capillary column (5 m, 0.32 mm i.d., 0.50 µm film thickness; Phenomenex, England), as previously described (Podechard et al., 2018). The oven temperature was programmed to increase from 200 to 350°C at a rate of 5°C/min, and the carrier gas was hydrogen (0.5 bar). The injector and the detector were set to 315°C and 345°C, respectively.

### Histology

Paraformaldehyde-fixed, paraffin-embedded liver tissue was sliced into 3 µm sections and stained with hematoxylin and eosin (H&E) or into 5 µm sections and stained with Sirius red or used for immuno-histochemistry. The staining was visualized with a light microscopy equipped with a Leica DFC300 camera. All liver sections were analyzed blindly. Liver steatosis was evaluated according to Kleiner (Kleiner et al., 2005). Steatosis was measured depending on the percentage of hepatocytes containing fat, where Grade 0 = less of 5% of hepatocytes containing fat in any section; grade 1 = 5% to 32% of hepatocytes; grade 2 = 33% to 65% of hepatocytes; grade 3 = up to 65% of hepatocytes. The degree of inflammation was appreciated by counting the inflammatory foci into 10 distinct areas at 200X for each liver slice (grade 0 = no foci; grade 1 = less of 2 foci per 200X field; grade 2 = 2 to 4 foci per 200X field; grade 3 = up to 4 foci per 200X field). Sirius Red staining was used for evaluation of fibrosis in the liver. Briefly, paraffin-embedded liver blocks (5 µm) sections were incubated with picrosirius red (#ab150681, Abcam, France) solution for 1h and quickly washed with acidified water (0.5% acetic acid). Collagen was selectively visualized as bright orange-red birefringent fibers and the image acquired under polarized light microscope. The fibrosis score was defined as follows: 0, no fibrosis; 1, pericellular and perivenular fibrosis; and 2, focal bridging fibrosis. Immunohistochemical staining of anti α-smooth muscle actin (α-SMA) (Abcam®) was also performed. Then, the peroxidase activity was revealed with diaminobenzidine (DAB, Dako K3468) and sections were counterstained rapidly with hematoxylin. α-SMA labelling score was established according to Akpolat (Akpolat et al., 2005) as follows; score 0: absence of labelling or <3% of periportal region; 1: between 3 and 33%; 2: between 34 and 66% and 3:>67% of the periportal region.

Lipid droplets detection, measurement and distribution visualization was performed using Python Jupyter Notebooks. Lipid droplets were detected using Cellpose with a model trained for lipid droplet segmentation (Stringer et al., 2021), applied at different scales (6.8 and 20 µm). The resulting label images are then filtered with a size bandwidth adapted to each scale. Then these different scales are merged by adding them successively in increasing order. Labels were added if less than 10% of their area is occupied by other labels from smaller scales. On the resulting multi- scale image labels area are measured, then converted to an equivalent diameter. From these measurements, LD diameter distribution was estimated for each tissue with a weighted kernel density estimation using the seaborn library (Waskom, 2021). The weight used was the area of each detection.

### Gene expression

Total cellular RNA from liver and BAT was extracted with Trizol Reagent® (Molecular Research Center, Inc., Cincinnati, OH, USA). RNAs were quantified using a nanophotometer (Implen). Total RNA samples (2 μg) were then reverse transcribed using the High-Capacity cDNA Reverse Transcription kit (Applied Biosystems^TM^) for real-time quantitative polymerase chain reaction (qPCR) analyses. The primers used for the SYBR Green assays are presented in Supplementary (Table S5). Amplifications were performed on a Stratagene Mx3005P thermocycler (Agilent Technology, Santa Clara, CA, USA). The qPCR data were normalized to the level of TATA-box binding protein (TBP) messenger RNA (mRNA), and analysed with LinRegPCR software to determine mean efficiency (NO), which was calculated as follows: NO = threshold/(Eff meanCq), where Eff mean: mean PCR efficiency, and Cq: quantification cycle.

Microarray experiments were conducted on n = 7 mice per group (3 or 4 per cage). Gene expression profiles were performed at the GeT-TRiX facility (GénoToul, Génopole Toulouse Midi-Pyrénées, France) using Sureprint G3 Mouse GE v2 microarrays (8 x 60K, design 074809, Agilent Technologies) following the manufacturer’s instructions. For each sample, Cyanine-3 (Cy3) labelled cRNA was prepared from 200bng of total RNA using the One-Color Quick Amp Labeling kit (Agilent technology) according to the manufacturer’s instructions. Then purification was performed by Agencourt RNAClean XP (Agencourt Bioscience Corporation, Beverly, Massachusetts, USA). Dye incorporation and cRNA yield were checked using Dropsense™ 96 UV/VIS droplet reader (Trinean, Gent, Belgium). A total of 600bng of Cy3-labelled cRNA were hybridized on the microarray slides following the manufacturer’s instructions. Immediately after washing, the slides were scanned on Agilent G2505C Microarray Scanner using Agilent Scan Control A.8.5.1 software. The fluorescence signal was extracted using Agilent Feature Extraction software v10.10.1.1 with default parameters. All experimental details and microarray data are available in NCBI’s Gene Expression Omnibus (Edgar et al., 2002) and are accessible through Gene Expression Omnibus (GEO: GSE279895; GSE279897; GSE280075).

### Quantification and Statistical analysis

All data are presented as means ± standard error of the mean (SEM). Statistical analysis on biochemical and qPCR data were performed using GraphPad Prism version 9 for Windows (GraphPad Software, San Diego, CA). One-way ANOVA was performed followed by appropriate post-hoc tests (Sidak’s multiple comparisons test) when differences were found to be significant (p<0.05). For histological scores, non-parametric test (Kruskall-Wallis) were used. Significance were indicated by: * or ^#^ for p < 0.05, ** or ^##^ for p < 0.01, *** or ^###^ for p < 0.001, **** or ^####^ for p < 0.0001.

Microarray data analyses were performed using R (R Core Team, 2018) and Bioconductor packages (Huber et al., 2015), as described in GSE accession GSE279895; GSE279897; GSE280075. Raw data (median signal intensity) were filtered, log2 transformed and normalized using quantile method qsmooth method (Hicks et al., 2018). A model was fitted using the limma ImFit function (Ritchie et al., 2015). Pair-wise comparisons between biological conditions were applied using specific contrasts. A correction for multiple testing was applied using the Benjamini-Hochberg procedure (BH), (Benjamini and Hochberg, 1995) to control the False Discovery Rate (FDR). Probes with FDR ≤0.05 were considered to be differentially expressed between conditions. Hierarchical clustering was applied to the samples and the differentially expressed probes using 1-Pearson correlation coefficient as distance and Ward’s criterion for agglomeration. The clustering results are illustrated as a heatmap of expression signals. The enrichment of Gene Ontology (GO) Biological Processes was evaluated using Metascape functions (Zhou et al., 2019).

## Data and code availability

- Microarray data have been deposited at GEO and are publicly available as of the date of publication. Accession numbers are listed in the Key resources table. All data reported in this paper will be shared by the lead contact upon request.
- This paper does not report original code.
- Any additional information required to reanalyze the data reported in this paper is available from the lead contact upon request.

## ACKNOWLEDGEMENTS

This work was supported by the French Foundation for the Medical Research FRM (Equipe FRM EQU202303016327). This work was also supported by grant from the French National Research Agency (ANR) IMAGINE (ANR-20-CE14-0038) and the Hepatomics FEDER program of Région Occitanie. We thank Anexplo (Genotoul, Toulouse) for their excellent work on plasma biochemistry.

## AUTHOR CONTRIBUTIONS

Conceptualization: N.L., A.F., and H.G.; Formal analysis: C.M.P.M., V.A.B., A.P., Y.L., J.B.M., E.M., B.C., S.L., T.L., H.G., S.E.S., A.F., and N.L.; Investigation: C.M.P.M., V.A.B., A.P., F.L., F.B., C.N., M.H., C.R., J.B., V.M., J.H.W., J.B.M., Y.L., E.R.B., E.M., R.F.F., C.D., B.C., S.L., D.L., T.L., L.G.P., H.G., S.E.S., A.F., and N.L.; Writing—original draft preparation, C.M.P.M., S.E.S., A.F., and N.L.; Writing—review and editing, S.E.S, N.L., A.F., and H.G.; Visualization, C.M.P.M., A.P., S.E.S., and N.L.; Supervision, S.E.S, N.L., and H.G.; Funding acquisition, N.L., S.E.S., and H.G. All authors have read and agreed to the published version of the manuscript.

## DECLARATION OF INTERESTS

The authors declare no competing interests.

### Supplemental information Legends

Document S1. Figures S1–S4 and Table S2, Table S5

## Notes

### Competing Interest Statement

The authors have declared no competing interest.

### Summary of Updates

typo correction and supplemental files added

## REFERENCES

Akpolat, N., Yahsi, S., Godekmerdan, A., Yalniz, M., and Demirbag, K. (2005). The value of alpha- SMA in the evaluation of hepatic fibrosis severity in hepatitis B infection and cirrhosis development: a histopathological and immunohistochemical study. Histopathology 47, 276–280.

Badman, M.K., Pissios, P., Kennedy, A.R., Koukos, G., Flier, J.S., and Maratos-Flier, E. (2007). Hepatic fibroblast growth factor 21 is regulated by PPARalpha and is a key mediator of hepatic lipid metabolism in ketotic states. Cell Metab. 5, 426–437.

Benjamini, Y., and Hochberg, Y. (1995). Controlling the False Discovery Rate: A Practical and Powerful Approach to Multiple Testing. J. R. Stat. Soc. Ser. B 57, 289–300.

Bligh, E.G., and Dyer, W.J. (1959). A rapid method of total lipid extraction and purification. Can. J. Biochem. Physiol. 37, 911–917.

Blondin, D.P., Frisch, F., Phoenix, S., Guérin, B., Turcotte, É.E., Haman, F., Richard, D., and Carpentier, A.C. (2017). Inhibition of Intracellular Triglyceride Lipolysis Suppresses Cold-Induced Brown Adipose Tissue Metabolism and Increases Shivering in Humans. Cell Metab. 25, 438–447.

Bucsek, M.J., Qiao, G., MacDonald, C.R., Giridharan, T., Evans, L., Niedzwecki, B., Liu, H., Kokolus, K.M., Eng, J.W.L., Messmer, M.N., et al. (2017). β-Adrenergic Signaling in Mice Housed at Standard Temperatures Suppresses an Effector Phenotype in CD8+ T Cells and Undermines Checkpoint Inhibitor Therapy. Cancer Res. 77, 5639–5651.

Cypess, A.M., Weiner, L.S., Roberts-Toler, C., Elía, E.F., Kessler, S.H., Kahn, P.A., English, J., Chatman, K., Trauger, S.A., Doria, A., et al. (2015). Activation of human brown adipose tissue by a β3-adrenergic receptor agonist. Cell Metab. 21, 33–38.

Donnelly, K.L., Smith, C.I., Schwarzenberg, S.J., Jessurun, J., Boldt, M.D., and Parks, E.J. (2005). Sources of fatty acids stored in liver and secreted via lipoproteins in patients with nonalcoholic fatty liver disease. J. Clin. Invest. 115, 1343–1351.

Edgar, R., Domrachev, M., and Lash, A.E. (2002). Gene Expression Omnibus: NCBI gene expression and hybridization array data repository. Nucleic Acids Res. 30, 207–210.

Eng, J.W.L., Reed, C.B., Kokolus, K.M., Pitoniak, R., Utley, A., Bucsek, M.J., Ma, W.W., Repasky, E.A., and Hylander, B.L. (2015). Housing temperature-induced stress drives therapeutic resistance in murine tumour models through β2-adrenergic receptor activation. Nat. Commun. 6.

Eslam, M., Newsome, P.N., Sarin, S.K., Anstee, Q.M., Targher, G., Romero-Gomez, M., Zelber-Sagi, S., Wai-Sun Wong, V., Dufour, J.F., Schattenberg, J.M., et al. (2020a). A new definition for metabolic dysfunction-associated fatty liver disease: An international expert consensus statement. J. Hepatol. 73, 202–209.

Eslam, M., Sanyal, A.J., George, J., Sanyal, A., Neuschwander-Tetri, B., Tiribelli, C., Kleiner, D.E., Brunt, E., Bugianesi, E., Yki-Järvinen, H., et al. (2020b). MAFLD: A Consensus-Driven Proposed Nomenclature for Metabolic Associated Fatty Liver Disease. Gastroenterology 158, 1999–2014.e1.

Fabbrini, E., Mohammed, B.S., Magkos, F., Korenblat, K.M., Patterson, B.W., and Klein, S. (2008). Alterations in adipose tissue and hepatic lipid kinetics in obese men and women with nonalcoholic fatty liver disease. Gastroenterology 134, 424–431.

Feldmann, H.M., Golozoubova, V., Cannon, B., and Nedergaard, J. (2009). UCP1 ablation induces obesity and abolishes diet-induced thermogenesis in mice exempt from thermal stress by living at thermoneutrality. Cell Metab. 9, 203–209.

Fischer, A.W., Cannon, B., and Nedergaard, J. (2018). Optimal housing temperatures for mice to mimic the thermal environment of humans: An experimental study. Mol. Metab. 7, 161–170.

Fougerat, A., Schoiswohl, G., Polizzi, A., Régnier, M., Wagner, C., Smati, S., Fougeray, T., Lippi, Y., Lasserre, F., Raho, I., et al. (2022). ATGL-dependent white adipose tissue lipolysis controls hepatocyte PPARα activity. Cell Rep. 39.

Gallage, S., Avila, J.E.B., Ramadori, P., Focaccia, E., Rahbari, M., Ali, A., Malek, N.P., Anstee, Q.M., and Heikenwalder, M. (2022). A researcher’s guide to preclinical mouse NASH models. Nat. Metab. 4.

Ganeshan, K., and Chawla, A. (2017). Warming the mouse to model human diseases. Nat. Rev. Endocrinol. 13, 458–465.

Giles, D.A., Ramkhelawon, B., Donelan, E.M., Stankiewicz, T.E., Hutchison, S.B., Mukherjee, R., Cappelletti, M., Karns, R., Karp, C.L., Moore, K.J., et al. (2016). Modulation of ambient temperature promotes inflammation and initiates atherosclerosis in wild type C57BL/6 mice. Mol. Metab. 5, 1121–1130.

Giles, D.A., Moreno-Fernandez, M.E., Stankiewicz, T.E., Graspeuntner, S., Cappelletti, M., Wu, D., Mukherjee, R., Chan, C.C., Lawson, M.J., Klarquist, J., et al. (2017). Thermoneutral housing exacerbates nonalcoholic fatty liver disease in mice and allows for sex-independent disease modeling. Nat. Med. 23, 829–838.

Haemmerle, G., Lass, A., Zimmermann, R., Gorkiewicz, G., Meyer, C., Rozman, J., Heldmaier, G., Maier, R., Theussl, C., Eder, S., et al. (2006). Defective lipolysis and altered energy metabolism in mice lacking adipose triglyceride lipase. Science 312, 734–737.

Hankir, M.K., and Klingenspor, M. (2018). Brown adipocyte glucose metabolism: a heated subject. EMBO Rep. 19.

Heine, M., Fischer, A.W., Schlein, C., Jung, C., Straub, L.G., Gottschling, K., Mangels, N., Yuan, Y., Nilsson, S.K., Liebscher, G., et al. (2018). Lipolysis Triggers a Systemic Insulin Response Essential for Efficient Energy Replenishment of Activated Brown Adipose Tissue in Mice. Cell Metab. 28, 644–655.e4.

Hicks, S.C., Okrah, K., Paulson, J.N., Quackenbush, J., Irizarry, R.A., and Bravo, H.C. (2018). Smooth quantile normalization. Biostatistics 19, 185–198.

Horakova, O., Sistilli, G., Kalendova, V., Bardova, K., Mitrovic, M., Cajka, T., Irodenko, I., Janovska, P., Lackner, K., Kopecky, J., et al. (2023). Thermoneutral housing promotes hepatic steatosis in standard diet-fed C57BL/6N mice, with a less pronounced effect on NAFLD progression upon high- fat feeding. Front. Endocrinol. (Lausanne). 14.

Huber, W., Carey, V.J., Gentleman, R., Anders, S., Carlson, M., Carvalho, B.S., Bravo, H.C., Davis, S., Gatto, L., Girke, T., et al. (2015). Orchestrating high-throughput genomic analysis with Bioconductor. Nat. Methods 12, 115–121.

Inagaki, T., Dutchak, P., Zhao, G., Ding, X., Gautron, L., Parameswara, V., Li, Y., Goetz, R., Mohammadi, M., Esser, V., et al. (2007). Endocrine regulation of the fasting response by PPARalpha-mediated induction of fibroblast growth factor 21. Cell Metab. 5, 415–425.

Kersten, S., Seydoux, J., Peters, J.M., Gonzalez, F.J., Desvergne, B., and Wahli, W. (1999). Peroxisome proliferator-activated receptor alpha mediates the adaptive response to fasting. J. Clin. Invest. 103, 1489–1498.

Kleiner, D.E., Brunt, E.M., Natta, M. Van, Behling, C., Contos, M.J., Cummings, O.W., Ferrell, L.D., Liu, Y.-C., Torbenson, M.S., Unalp-Arida, A., et al. (2005). Design and Validation of a Histological Scoring System for Nonalcoholic Fatty Liver Disease.

Kokolus, K.M., Capitano, M.L., Lee, C.T., Eng, J.W.L., Waight, J.D., Hylander, B.L., Sexton, S., Hong, C.C., Gordon, C.J., Abrams, S.I., et al. (2013). Baseline tumor growth and immune control in laboratory mice are significantly influenced by subthermoneutral housing temperature. Proc. Natl. Acad. Sci. U. S. A. 110, 20176–20181.

Labbé, S.M., Caron, A., Bakan, I., Laplante, M., Carpentier, A.C., Lecomte, R., and Richard, D. (2015). In vivo measurement of energy substrate contribution to cold-induced brown adipose tissue thermogenesis. FASEB J. 29, 2046–2058.

Lee, E., Korf, H., and Vidal-Puig, A. (2023). An adipocentric perspective on the development and progression of non-alcoholic fatty liver disease. J. Hepatol. 78, 1048–1062.

Li, Y., Fromme, T., Schweizer, S., Schöttl, T., and Klingenspor, M. (2014). Taking control over intracellular fatty acid levels is essential for the analysis of thermogenic function in cultured primary brown and brite/beige adipocytes. EMBO Rep. 15, 1069–1076.

Low, Z.S., Chua, D., Cheng, H.S., Tee, R., Tan, W.R., Ball, C., Sahib, N.B.E., Ng, S.S., Qu, J., Liu, Y., et al. (2024). The LIDPAD Mouse Model Captures the Multisystem Interactions and Extrahepatic Complications in MASLD. Adv. Sci. (Weinheim, Baden-Wurttemberg, Ger.

Lu, D., He, A., Tan, M., Mrad, M., El Daibani, A., Hu, D., Liu, X., Kleiboeker, B., Che, T., Hsu, F.F., et al. (2024). Liver ACOX1 regulates levels of circulating lipids that promote metabolic health through adipose remodeling. Nat. Commun. 15.

Maloney, S.K., Fuller, A., Mitchell, D., Gordon, C., and Michael Overton, J. (2014). Translating animal model research: does it matter that our rodents are cold? Physiology (Bethesda). 29, 413– 420.

Marcher, A.B., Loft, A., Nielsen, R., Vihervaara, T., Madsen, J.G.S., Sysi-Aho, M., Ekroos, K., and Mandrup, S. (2015). RNA-Seq and Mass-Spectrometry-Based Lipidomics Reveal Extensive Changes of Glycerolipid Pathways in Brown Adipose Tissue in Response to Cold. Cell Rep. 13, 2000–2013.

Montagner, A., Polizzi, A., Fouché, E., Ducheix, S., Lippi, Y., Lasserre, F., Barquissau, V., Régnier, M., Lukowicz, C., Benhamed, F., et al. (2016). Liver PPARα is crucial for whole-body fatty acid homeostasis and is protective against NAFLD. Gut 65, 1202–1214.

Morrow, M.R., Batchuluun, B., Wu, J., Ahmadi, E., Leroux, J.M., Mohammadi-Shemirani, P., Desjardins, E.M., Wang, Z., Tsakiridis, E.E., Lavoie, D.C.T., et al. (2022). Inhibition of ATP-citrate lyase improves NASH, liver fibrosis, and dyslipidemia. Cell Metab. 34, 919–936.e8.

Nunes, J.R.C., Smith, T.K.T., Ghorbani, P., O’Dwyer, C., Trzaskalski, N.A., Dergham, H., Pember, C., Kilgour, M.K., Mulvihill, E.E., and Fullerton, M.D. (2023). Thermoneutral housing does not accelerate metabolic dysfunction-associated fatty liver disease in male or female C57Bl/6J mice fed a Western diet. Am. J. Physiol. Endocrinol. Metab. 325, E10–E20.

Oates, J.R., Sawada, K., Giles, D.A., Alarcon, P.C., Damen, M.S.M.A., Szabo, S., Stankiewicz, T.E., Moreno-Fernandez, M.E., and Divanovic, S. (2023). Thermoneutral housing shapes hepatic inflammation and damage in mouse models of non-alcoholic fatty liver disease. Front. Immunol. 14.

Orava, J., Nuutila, P., Lidell, M.E., Oikonen, V., Noponen, T., Viljanen, T., Scheinin, M., Taittonen, M., Niemi, T., Enerbäck, S., et al. (2011). Different metabolic responses of human brown adipose tissue to activation by cold and insulin. Cell Metab. 14, 272–279.

Parlati, L., Régnier, M., Guillou, H., and Postic, C. (2021). New targets for NAFLD. JHEP Reports Innov. Hepatol. 3.

Podechard, N., Ducheix, S., Polizzi, A., Lasserre, F., Montagner, A., Legagneux, V., Fouché, E., Saez, F., Lobaccaro, J.M., Lakhal, L., et al. (2018). Dual extraction of mRNA and lipids from a single biological sample. Sci. Rep. 8.

Régnier, M., Polizzi, A., Lippi, Y., Fouché, E., Michel, G., Lukowicz, C., Smati, S., Marrot, A., Lasserre, F., Naylies, C., et al. (2018). Insights into the role of hepatocyte PPARα activity in response to fasting. Mol. Cell. Endocrinol. 471, 75–88.

Riazi, K., Azhari, H., Charette, J.H., Underwood, F.E., King, J.A., Afshar, E.E., Swain, M.G., Congly, S.E., Kaplan, G.G., and Shaheen, A.A. (2022). The prevalence and incidence of NAFLD worldwide: a systematic review and meta-analysis. Lancet. Gastroenterol. Hepatol. 7, 851–861.

Rinella, M.E., Neuschwander-Tetri, B.A., Siddiqui, M.S., Abdelmalek, M.F., Caldwell, S., Barb, D., Kleiner, D.E., and Loomba, R. (2023). AASLD Practice Guidance on the clinical assessment and management of nonalcoholic fatty liver disease. Hepatology 77, 1797–1835.

Ritchie, M.E., Phipson, B., Wu, D., Hu, Y., Law, C.W., Shi, W., and Smyth, G.K. (2015). limma powers differential expression analyses for RNA-sequencing and microarray studies. Nucleic Acids Res. 43, e47.

Roberts-Toler, C., O’Neill, B.T., and Cypess, A.M. (2015). Diet-induced obesity causes insulin resistance in mouse brown adipose tissue. Obesity (Silver Spring). 23, 1765–1770.

Rosen, E.D., and Spiegelman, B.M. (2006). Adipocytes as regulators of energy balance and glucose homeostasis. Nature 444, 847–853.

Schreiber, R., Diwoky, C., Schoiswohl, G., Feiler, U., Wongsiriroj, N., Abdellatif, M., Kolb, D., Hoeks, J., Kershaw, E.E., Sedej, S., et al. (2017). Cold-Induced Thermogenesis Depends on ATGL-Mediated Lipolysis in Cardiac Muscle, but Not Brown Adipose Tissue. Cell Metab. 26, 753–763.e7.

Seeley, R.J., and MacDougald, O.A. (2021). Mice as experimental models for human physiology: when several degrees in housing temperature matter. Nat. Metab. 3, 443–445.

Shin, H., Ma, Y., Chanturiya, T., Cao, Q., Wang, Y., Kadegowda, A.K.G., Jackson, R., Rumore, D., Xue, B., Shi, H., et al. (2017). Lipolysis in Brown Adipocytes Is Not Essential for Cold-Induced Thermogenesis in Mice. Cell Metab. 26, 764–777.e5.

Simcox, J., Geoghegan, G., Maschek, J.A., Bensard, C.L., Pasquali, M., Miao, R., Lee, S., Jiang, L., Huck, I., Kershaw, E.E., et al. (2017). Global Analysis of Plasma Lipids Identifies Liver-Derived Acylcarnitines as a Fuel Source for Brown Fat Thermogenesis. Cell Metab. 26, 509–522.e6.

Škop, V., Guo, J., Liu, N., Xiao, C., Hall, K.D., Gavrilova, O., and Reitman, M.L. (2020). Mouse Thermoregulation: Introducing the Concept of the Thermoneutral Point. Cell Rep. 31.

Škop, V., Xiao, C., Liu, N., Gavrilova, O., and Reitman, M.L. (2021). The effects of housing density on mouse thermal physiology depend on sex and ambient temperature. Mol. Metab. 53.

Smati, S., Polizzi, A., Fougerat, A., Ellero-Simatos, S., Blum, Y., Lippi, Y., Régnier, M., Laroyenne, A., Huillet, M., Arif, M., et al. (2022). Integrative study of diet-induced mouse models of NAFLD identifies PPARα as a sexually dimorphic drug target. Gut 71, 807–821.

Stringer, C., Wang, T., Michaelos, M., and Pachitariu, M. (2021). Cellpose: a generalist algorithm for cellular segmentation. Nat. Methods 18, 100–106.

Tian, X.Y., Ganeshan, K., Hong, C., Nguyen, K.D., Qiu, Y., Kim, J., Tangirala, R.K., Tonotonoz, P., and Chawla, A. (2016). Thermoneutral Housing Accelerates Metabolic Inflammation to Potentiate Atherosclerosis but Not Insulin Resistance. Cell Metab. 23, 165–178.

Vacca, M., Kamzolas, I., Harder, L.M., Oakley, F., Trautwein, C., Hatting, M., Ross, T., Bernardo, B., Oldenburger, A., Hjuler, S.T., et al. (2024). An unbiased ranking of murine dietary models based on their proximity to human metabolic dysfunction-associated steatotic liver disease (MASLD). Nat. Metab. 6, 1178–1196.

Vialard, F., and Olivier, M. (2020). Thermoneutrality and Immunity: How Does Cold Stress Affect Disease? Front. Immunol. 11.

Waskom, M.L. (2021). seaborn: statistical data visualization. J. Open Source Softw. 6, 3021.

X, L., M, R., J, M., M, R., ME, H., and LP, K. (2003). Paradoxical resistance to diet-induced obesity in UCP1-deficient mice. J. Clin. Invest. 111, 399–407.

Younossi, Z.M., Golabi, P., de Avila, L., Paik, J.M., Srishord, M., Fukui, N., Qiu, Y., Burns, L., Afendy, A., and Nader, F. (2019). The global epidemiology of NAFLD and NASH in patients with type 2 diabetes: A systematic review and meta-analysis. J. Hepatol. 71, 793–801.

Yu, X.X., Lewin, D.A., Forrest, W., and Adams, S.H. (2002). Cold elicits the simultaneous induction of fatty acid synthesis and beta-oxidation in murine brown adipose tissue: prediction from differential gene expression and confirmation in vivo. FASEB J. 16, 155–168.

Zhang, S., Williams, K.J., Verlande-Ferrero, A., Chan, A.P., Su, G.B., Kershaw, E.E., Cox, J.E., Maschek, J.A., Shapira, S.N., Christofk, H.R., et al. (2024). Acute activation of adipocyte lipolysis reveals dynamic lipid remodeling of the hepatic lipidome. J. Lipid Res. 65.

Zhou, X.-D., Lonardo, A., Pan, C.Q., Shapiro, M.D., Zheng, M.-H., Zheng, K.I., Ma, H.-L., Zhu, P.-W., Pan, X.-Y., Zhang, R., et al. (2024). Clinical Features and Long-Term Outcomes of Patients Diagnosed with MASLD, MAFLD, or Both. J. Hepatol.

Zhou, Y., Zhou, B., Pache, L., Chang, M., Khodabakhshi, A.H., Tanaseichuk, O., Benner, C., and Chanda, S.K. (2019). Metascape provides a biologist-oriented resource for the analysis of systems-level datasets. Nat. Commun. 10.

