## Supplemental Figures, Table S2, Table S5 for "Effect of thermoneutral housing on MASLD severity, hepatic gene expression, and BAT activation during β3-adrenergic stimulation in mice"

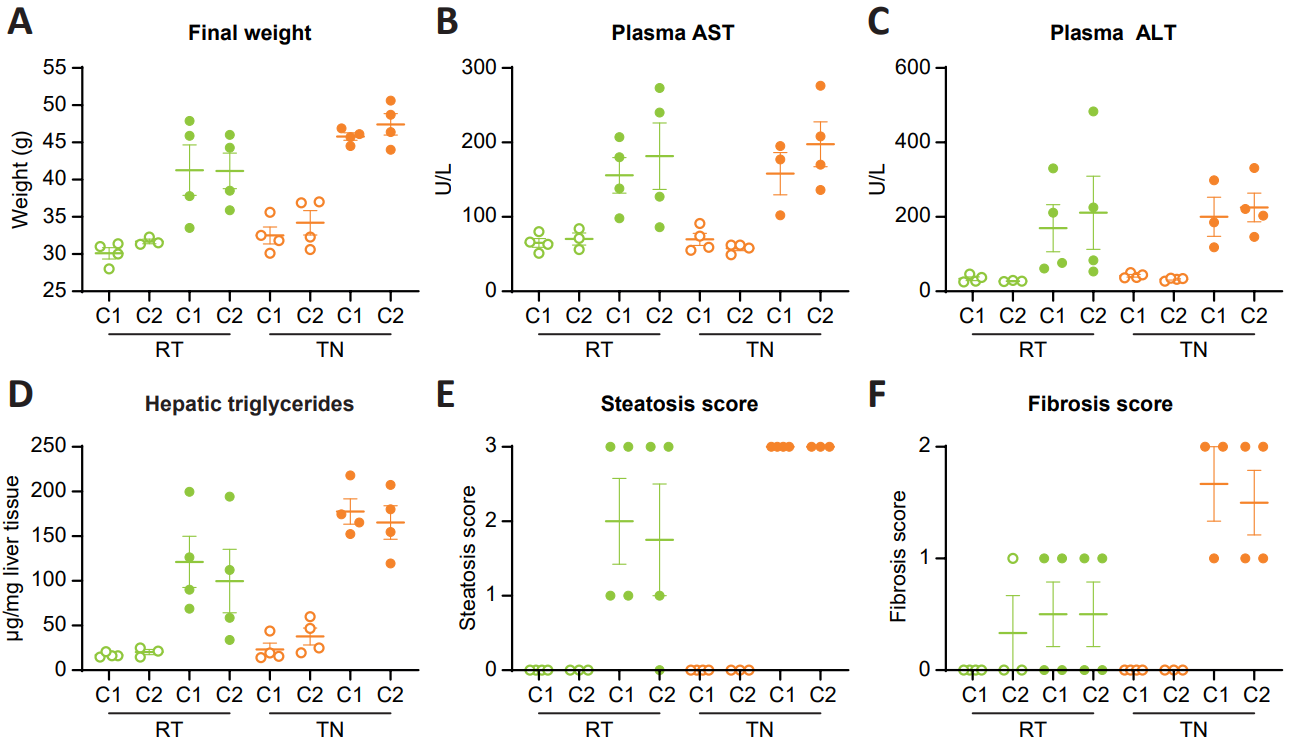

**Supplementary Figure S1: Thermoneutral housing reduces the intra-group variability in WD-induced MASLD.** (**A**) Final weight after 13 weeks of diet in each cage (C1 or C2) under room temperature (RT) or thermoneutral housing (TN). (**B, C**) Plasma ALT and AST activity. ALT alanine aminotransferase, AST aspartate aminotransferase. (**D**) Hepatic triglycerides extracted from the livers and analysed by gas-liquid chromatography. (**E**) Liver steatosis estimated on histological liver sections. Scoring: parenchymal involvement by steatosis <5%, 0; 5%-33%, 1; 33%-66%, 2; >66%, 3 (n=4/group). (**F**) Stellate hepatic cell activation estimated on histological liver sections. Scoring: lower than 3% of periportal area, 0; 3%-33%, 1; 34%-66%, 2; more than 66%, 3 (n=4/group).

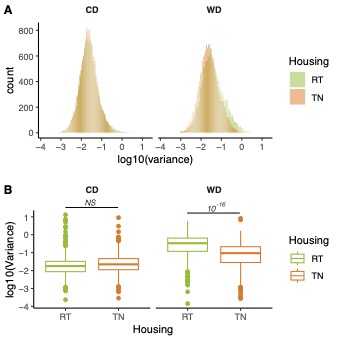

**Supplementary Figure S2: Thermoneutral housing reduces the intra-group variability in WD-induced changes in hepatic gene expression.** (**A**) Distribution of the variances per experimental group for all hepatic genes measured using microarrays. (**B**) Variances of the 449 non-homoscedastic hepatic genes.

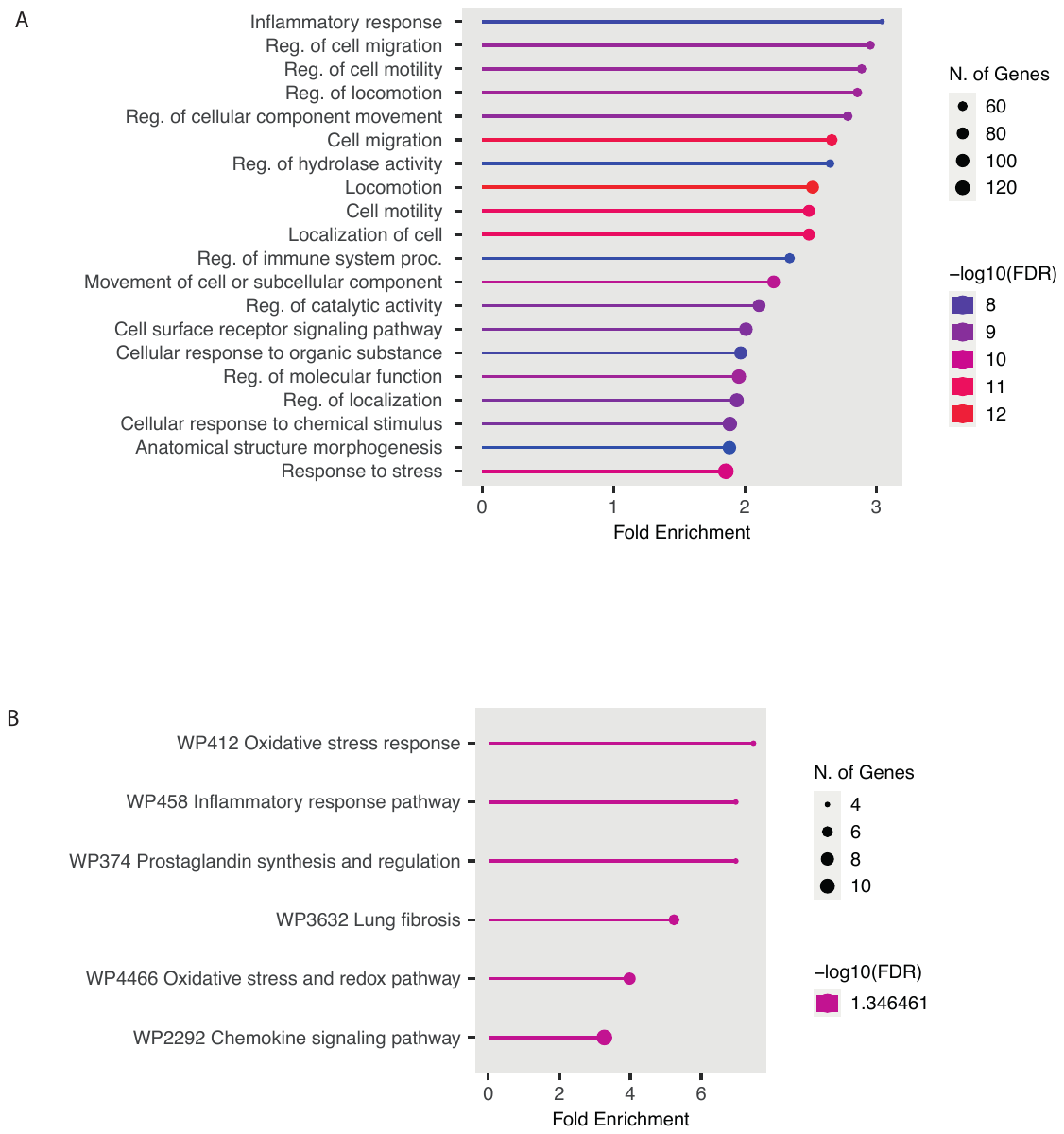

**Supplementary Figure S3: Thermoneutral housing reduces the intra-group variability in WD-induced changes in hepatic gene expression.** The most significantly enriched biological processes identified with the Metascape gene ontology algorithm are shown at the right of each profile. Briefly, hypergeometric tests were performed for each category in each cluster. (**A**) Distribution of the variances per experimental group for all hepatic genes measured using microarrays. (**B**) Variances of the 449 non-homoscedastic hepatic genes.

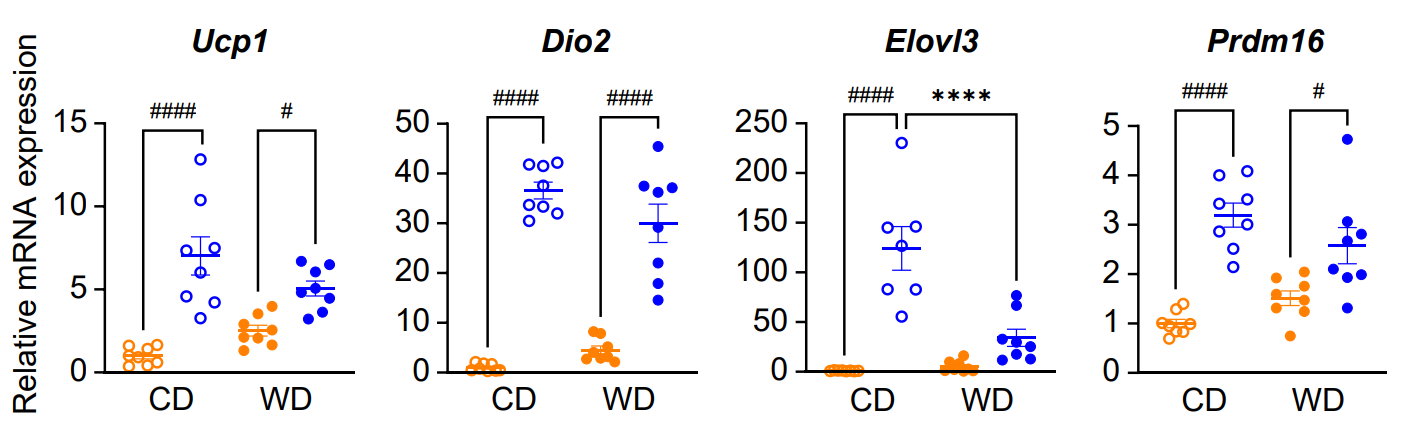

**Supplementary Figure S4: β3-adrenergic signalling induced-BAT activation is blunted in MASLD condition.** Related to Figure 5. mRNA relative expression of *Ucp1*, *Dio2*, *Elovl3* and *Prdm16* measured by qRT-PCR in the brown adipose tissue (BAT). Data are presented as the mean ± SEM for n = 8/group. * diet effect, # CL316243 effect. * or #P < 0,05, **P < 0,01, *** or ###P < 0,001, **** or ####P < 0,0001. Differential effects were analysed by analysis of variance (one-way ANOVA) with post hoc Šídák’s test.

**Supplementary Table S2:** Descriptive statistics for selected metabolic parameters.

| **Parameter** |  | **CD RT** | **WD RT** | **CD TN** | **WD TN** |
| --- | --- | --- | --- | --- | --- |
| Body weight (g) | Mean | 30.8 | 41.2 | 33.3 | 46.6 |
|  | SD | 1.4 | 5.4 | 2.7 | 2.1 |
|  | SEM | 0.5 | **1.9** | 0.97 | **0.7** |
| Plasma ALT (U/L) | Mean | 31 | 190.3 | 36.9 | 214.6 |
|  | SD | 7.7 | 154.4 | 7.1 | 77 |
|  | SEM | 2.9 | **54.6** | 2.5 | **29.1** |
| Hepatic triglycerides (µg/mg tissue | Mean | 18.3 | 110.5 | 30.5 | 171.5 |
|  | SD | 3.9 | 60.9 | 17.2 | 31.5 |
|  | SEM | 1.5 | **21.5** | 6.1 | **11.1** |

**Supplementary Table S5:** Oligonucleotide sequences for real-time qPCR.

**Related to STAR methods**

| **Gene** | **NCBI RefSeq** | **Forward primer** | **Reverse primer** |
| --- | --- | --- | --- |
| *Acta2* | NM_007392 | ACCCACCCAGAGTGGAGAA | GCTGTGCTGTCTTCCTCTTCA |
| *Col1a1* | NM_007742 | CTCCGGCTCCTGCTCCTC | GGGTTTCCACGTCTCACCAT |
| *Col3a1* | NM_009930 | TGGCACAGCAGTCCAACGTA | CATAGGACTGACCAAGGTGGCT |
| *Dio2* | NM_010050.4 | AGCTTCCTCCTAGATGCCTACA | GGGAGCATCTTCACCCAGTTTA |
| *Elovl3* | NM_007703 | GCCTCTCATCCTCTGGTCCT | TGCCATAAACTTCCACATCCT |
| *Il-1β* | NM_008361.4 | AAAAAAGCCTCGTGCTGTCG | GTCGTTGCTTGGTTCTCCTTGT |
| *Mmp13* | NM_008607 | ACCTGGACAAGCAGTTCCAAAG | AGCTCATGGGCAGCAACAATAA |
| *Prdm16* | NM_027504.3 | AAGGCGAGGGCGAGGAA | CATATTATTTACAACGTCACCGTCACT |
| *Tnfα* | NM_013693 | TCCCCAAAGGGATGAGAAGTTC | GCGCTGGCTCAGCCACT |
| *Ucp1* | NM_009463.3 | CCTGCCTCTCTCGGAAACAA | TGTAGGCTGCCCAATGAACA |
